# Chromatin interaction-based annotation of distal cis-regulatory elements reveals highly dynamic promoter-enhancer interactions in lymphocyte development

**DOI:** 10.1101/2025.05.15.653418

**Authors:** Johanna Tingvall-Gustafsson, Christina T. Jensen, Jonas Ungerbäck, Mikael Sigvardsson

## Abstract

Stage- and lineage-specific gene expression patterns are controlled by a complex interplay between transcription factors, the epigenetic landscape, and the 3-dimensional (3D) structure of the DNA. The 3D structure allows for the formation of DNA loops that juxtaposition distal regulatory elements to the promoters, allowing for tight control of gene expression. These loops can span hundreds of thousands base pairs, making it challenging to link regulatory elements to the correct target genes using conventional proximity-based annotation methods. Using a novel tool to facilitate the exploration of complex gene regulatory networks based on chromosome configuration data in early lymphocytes, we show that lineage-specific transcription factors target regulatory elements that are annotated to both lineage-specific and broadly expressed genes. Targeted inactivation of a set of these genes revealed their importance for B-cell development. Several regulatory elements annotated to lineage-specific genes were also annotated to alternative promoters in a context dependent manner, revealing a highly complex interplay between promoters and DREs in early lymphocyte development. Investigating the chromatin configuration in EBF1 deficient pro-B cells identified this transcription factor as a key mediator in the establishment of the 3-D promoter enhancer landscape in early B-lymphocyte differentiation. These data highlight how efficient annotation procedures for linking distal regulatory elements to target genes provide valuable insights into gene regulatory networks.

## Introduction

Differentiation of hematopoietic stem cells into highly differentiated blood cells requires the orchestrated action of transcription factors (TFs), establishing gene regulatory networks (GRNs). The TF networks that control the separation of the B- and T-lymphoid lineages have been extensively studied, and several key regulators of fate determination have been identified. T-lineage cell fate is dependent upon the TFs TCF7, GATA3 and BCL11B, which act in a functional hierarchy to activate T-lineage-specific gene expression programs and establish a stable commitment to the T-lineage cell fate (1–5). In a similar manner, the TFs TCF3, EBF1 and PAX5 act in a functional hierarchy to establish B-cell fate in lymphoid progenitors (6–12). While several key regulators of lymphoid cell fate determination have been identified, it remains to be resolved how these factors regulate transcription from target genes to drive lineage commitment and dictate the cell fates of multipotent progenitor cells. Although the binding sites for TFs in the genome can be identified, most of these cannot easily be assigned to any specific target gene using conventional annotation approaches (6, 8, 10, 11, 13).

Regulatory elements make up around 6% of the human genome, and it has been estimated that each promoter can interact with on average 5–50 distal regulatory elements (DREs) (14, 15). While most of the regulatory elements that affect transcription in budding yeasts are located within a few hundred base-pairs (bp) of their target promoters (16), DREs located several kilobases (kb) from their promoter elements have been identified in Drosophila (17). These elements are, however, mainly acting on proximal genes (17). In mammals, it has been estimated that the average distance between promoters and DREs lies in the range of 100–500 kb (14, 18–20), and only 27%–60% of these DREs act on their most-proximal promoter (21–23). It has been suggested that the ability of DREs to act over such long distances reflects the fact that the genome is arranged in a highly organized and dynamic three-dimensional (3D) architecture (24, 25). This architecture comprises large-scale structures of chromosomal territories, compartments and topologically associated domains (TADs), as well as the smaller-scale structures of chromatin loops (26, 27). These structures can serve to connect physically various DREs, such as enhancer elements, to promoters (28, 29), and it has been proposed that they are important for functional gene regulation (30–33). Even though the classical model for static promoter enhancer interactions is challenged (34–36), most models predict that the functional interplay between regulatory elements involves proximity in a 3D space at some point in the activation process (29).

Even though it is well established that DREs in human and mice act over long distances and not always at the most-proximal gene (21–23), the standard methods used for the annotation of DREs to target genes are based on proximity on the linear DNA strand. Improvements have been made to the standard proximity-based annotation to facilitate the annotation of more-distal peaks. This includes the use of the Genomic Regions Enrichment of Annotations Tool (GREAT) (37), which by introducing a concept of gene regulatory domains surrounding the transcription start site (TSS) and extending the annotation to domains of neighboring genes, allows for increased inter-element distances and the assignment of more than one gene for each DRE. However, all proximity-based methods are highly dependent upon the local gene density and are restricted by an upper distance limit for the identification of regulatory elements (Figure S1A-B). This local gene density dependence, combined with the inability to incorporate the dynamic aspect of chromatin organization into the annotation, highlights the need for improved annotation methods for DREs. As an alternative to proximity-based annotation, methods to link gene expression to DRE-accessibility has been employed (38). While this approach creates a functional link between a gene and a regulatory element, the method assumes that DRE activity is always reflected in changes in chromatin accessibility. With advancements in methods that capture chromatin-genome 3D organization (e.g., Hi-C, HiCHiP), interaction-based assignment of target gene to DREs has emerged as an alternative solution. Interaction-based annotation is not constrained by the simplistic linear representation of the genome and can incorporate cell type-specific or dynamic aspects of genome organization into the annotation process. There are also publicly available databases for chromatin interaction data (39), ensuring that interaction datasets are readily accessible for use in annotation. Despite the advantages of interaction-based methods, the use of chromatin interactions in the annotation of DREs remains limited, potentially related to the lack of use-friendly tools that doesn’t require extensive processing for incorporation of the interaction-data into the annotation process. To address the need for easy-to-use tools that can improve the annotation of DREs, we have generated an Interaction-based Cis-regulatory Element Annotator (ICE-A) (Figure S1C). ICE-A is based on the widely used and reproducibility-focused workflow management system Nextflow (40). It can, with a single command, perform interaction-based annotation of one or more sets of genomic regions (e.g., peak files from an ATAC-seq (Assay for Transposase-Accessible Chromatin using sequencing) or ChIP-seq experiment). In addition, ICE-A can perform interaction-centered annotation and provide several options for visualization of the results. Using this tool, we have investigated the regulatory landscapes in lymphoid progenitor cells. This has revealed that lineage-restricted TFs frequently bind to cis-regulatory elements that are annotated to broadly expressed genes, several of which are essential for B-lymphocyte development. Furthermore, regulatory elements annotated to lineage-specific genes could be annotated to alternative promoters in a context-dependent manner. The results show how interaction-based annotation of regulatory elements has the potential to increase significantly our understanding of GRN in health and disease.

## Methods

### EXPERIMENTAL MODEL AND STUDY PARTICIPANT DETAILS

#### Mice models

TetO-Cas9 mice (stock no: 029476 - B6.Cg-Col1a1<tm1(tetO-cas9)Sho>/J), bought from JAX, were crossed with R26m2rtTA mice (41) to obtain a Cas9 inducible (iCas9) mouse line. Both male and female mice aged 8-16 weeks were used for isolation of BM progenitor cells. Animal procedures were performed with consent from the local ethics committee at Lund University (Sweden).

#### Cell culturing

The DN3-line T cell progenitor cell line Scid.adh.2C2 is a subclone the Scid.adh cell line derived from a spontaneous thymic lymphoma in SCID mutant mouse (42). Scid.adh.2C2 T-cell progenitor cell line were cultured in RPMI1640 with 10% fetal bovine serum, sodium pyruvate, non-essential amino acids, gentamicin and 50 µM β-mercaptoethanol. The Abelson transformed pre-B cell-line 230-238 were maintained in RPMI1640 supplemented with 10% heat-inactivated fetal calf serum, 25mM HEPES, 50µg/ml gentamicin, and 50µM β-mercaptoethanol. Primary bone marrow cells were cultured on OP9 or OP9D stromal cells in OptiMEM media supplemented with 10% fetal calf serum, 50µg/mL gentamicin, 50µM β-mercaptoethanol, 50ng/mL KIT ligand, 50ng/mL Fms-like tyrosine kinase 3 ligand (FLT3L) and 50ng/mL interleukin-7 (IL-7). Cas9 expression was induced with 0.1 µg/mL Doxycyclin (DOX, Sigma). All cytokines were obtained from Peprotech.

### METHOD DETAILS

#### ICE-A

ICE-A is an open source tool for chromatin interaction-based annotation of genomic regions based on the widely used and reproducibility-focused workflow management system Nextflow (40). The tool allows for the assignment of target genes to one or more sets of genomic regions (bed format) using user-specified chromatin interactions. ICE-A accepts interactions in 2D-bed format with any number of columns, if the first six columns specify the genomic coordinates, making it compatible with different interaction-calling tools. ICE-A can be run on a standard laptop, with a typical run time for annotation of 4 peak files for basic and multiple mode is 2 and 8 min respectively. More information and installation instructions can be found at: https://github.com/Tingvall/ICE_A.

#### Interaction-based annotation

Interaction-based annotation of ICE-A is based on overlaps between the user-defined regions and interaction anchor points. A region overlapping an anchor point is assigned to a target gene if the corresponding interaction anchor overlaps with its promoter region (default: ±2,500 bp from the target TSS). Even though the main concept of ICE-A is to use chromatin interactions to improve the accuracy and specificity of the DRE annotation, the dependence on predefined interactions introduces constraints (e.g., bin size and minimum distance for interaction calling), thereby preventing the annotation of peaks to proximally located genes. Given the high likelihood that peaks located within promoter regions will influence the expression of the corresponding gene, disregarding these constraints would be inappropriate. To handle this aspect, ICE-A combines conventional proximity-based and interaction-based annotation systems to improve the accuracy and cell specificity of the annotation of distal regulatory regions, while still allowing for peaks that are located close to promoter regions to be annotated to the corresponding gene. For peaks located at a distance that is below the interaction threshold (default 2*bin size) to the closest TSS, the proximal annotation obtained from annotatePeaks.pl for HOMER (43) was applied, in addition to any available interaction-based annotations. The default behavior of ICE-A is to use only interactions that overlap with the peak in one of the anchor points and a promoter in the second anchor point. However, the resolution of the interaction-based annotation is limited by the bin size, with the consequence that loops where the true interaction is located several kilo-bases from the peak can be used for annotation, while closer interactions are ignored based on the arbitrarily defined bins. To make the interaction-based annotation less-stringent, ICE-A offers the possibility to also include neighboring interactions, defined by either the distance or number of bins from the bin overlapping the peak/promoter using the options: –close_peak_type and –close_promoter_type. Adjusting these parameters can also be suitable if the chromatin interactions used for annotation have been subjected to nearby interaction filtering, and in the case of an alternative TSS. As the default, ICE-A uses the TSS positions from HOMER, although it is possible to provide custom annotations to which the genomic regions can be assigned.

The main output from the peak-centered annotation is a single text file for each provided input bed with information about each annotation, including the target gene symbol, entrezID, distance to the target TSS, type of annotation used (proximal/interaction-based), and interaction score (if provided). In contrast to conventional proximity-based annotation, interaction-based annotation can generate multiple annotations per region. ICE-A offers several options for how to handle this depending on the intended subsequent analysis. The default behavior is to present all annotations in a comma-separated format with one row per peak. It is also possible to obtain the output file in a format with one row for each unique annotation. In addition to the annotated peak file(s), a gene list for each input bed file is provided, which is suitable for gene ontology analysis etc.

#### Modes

To account for different types of data, ICE-A can be run in three different modes. Basic mode performs interaction-based annotation of a single bed file. Multiple mode is suitable for cases where identification of overlaps between different sets of regions are of interest, e.g., co-occupancy of multiple transcriptional regulators. In addition to performing an annotation of every peak file, Multiple mode identifies overlaps defined either at the bin level or based on overlaps with customized regions (user-provided or a union of input bed files). Differential mode handles the common situation of comparing two conditions (e.g., differential TF binding). By providing the corresponding gene expression data, distally located peaks associated with changes in gene expression can be identified, as well as being categorized as activating or repressive.

#### Visualization

ICE-A provides numerous options for the visualization of peak annotations depending on the mode. If run in Multiple mode, the overlaps between a set of regions can be identified and visualized in the forms of UpSet and Circos plots. UpSet plots, which are based on the UpSetPlot library (a Python implementation of the UpSet plots (44)), show the overlaps of promoters and distally located regions, separately. For the Circos plot, overlaps between input bed files in interacting distal and proximal regions are visualized using the circlize R package (45). Both UpSet and Circos plots can be filtered on elements associated with a user-specified gene list if the option –filter_genes is used. For all three modes, there is the option to visualize the interaction in a network format using Cytoscape (46). If provided, interaction and/or peak scores can be represented by edge weight. If run in Differential mode, the network can be filtered according to differentially expressed genes, and separate networks for annotations associated with up- and down-regulated genes can be generated. In addition to the PDF output of the default network layout, ICE-A also provides the network xGMML files to be loaded into Cytoscape for visualization and customization in an interactive format.

#### Distance limits of proximity-based annotation

The theoretical distance limits for assignment of a distal element to a target gene were calculated for both the mouse (mm10) and human (hg38) genomes based on standard proximity annotation (annotation to the closest TSS) and GREAT. For standard proximity annotation, the upper threshold for assigning an element to a gene is half the distance to the upstream and down-stream neighboring genes, respectively. For GREAT, the distance is extended to the basal gene regulatory domain (default -5 kb/+1 kb from the TSS) of the neighboring genes. Curated regions provided by GREAT (v4.0) are included in the distance calculations. For both annotation methods, the TSS coordinates are based on the USCS Known Gene dataset (NCBI build 37 derived from Ensembl Biomart ver. 67 for mouse, and GRCh38 derived from Ensembl Biomart ver. 90 for human). A violin plot of the maximal distance for CRE annotation (irrespective of direction) for each gene, as well as a barplot of the median upper distance limit in the up- and down-stream direction, respectively, is generated using ggplot2 (v3.5.1).

#### ChIP-sequencing and data analysis

##### ChIP-seq

ChIP-seq was carried out as previously reported for histone modifications in Strid et al. (13) in duplicates. Scid.adh.2C T-cell progenitor cell line were fixed in 1% formaldehyde for 10 min at RT followed by quenching by addition of 1/10 volume of 0.125 M glycine and wash in PBS. Nuclei were isolated by incubation in Nuclei Isolation buffer (50 mM Tris, pH 8, 60 mM KCl, and 0.5% NP40) + protease inhibitor mixture (PIC) (Roche Diagnostics) for 10 min on ice. Pelleted nuclei were resuspended in Lysis buffer (0.5% SDS, 10 mM EDTA, 0.5 mM EGTA, and 50 mM Tris–HCl (pH 8))) + PIC and sonicated on a Bioruptor (Diagenode), followed by pelleting of debris. The supernatant was diluted 5◊ in HBSS (Lonza) + PIC and 2◊ radioimmunoprecipitation assay buffer (20 mM Tris–HCl (pH 7.5), 2 mM EDTA, 2% Triton X-100, 0.1% SDS, 0.2% sodium deoxycholate, and 200 mM NaCl) + PIC. 10µl of H3K4me3 (Millipore, 07-473) was hybridized to 70 µl Protein G/A Dynabeads (Life Technologies). ChIP was performed overnight at 4°C and subsequently washed 1x 500 µl Low Salt Immune Complex Wash Buffer (0.1% SDS, 1% Triton-X, 2 mM EDTA, 20 mM Tris-HCl (pH 8.1), 150 mM NaCl), 1x with 200 µl High Salt Immune Complex Wash Buffer, (0.1% SDS, 1% Triton-X, 2 mM EDTA, 20 mM Tris-HCl (pH 8.1), 500 mM NaCl), 1x with 200 µl LiCl Immune Complex Wash Buffer (0.25 M LiCl, 1% Igepal-CA630, 1% deoxycholic acid, 1 mM EDTA, 10 mM Tris (pH 8.1))), and 2x with 200 µl TE buffer (10 mM Tris-Hcl (pH 8.0), 10 mM EDTA). Chromatin was eluted for 6 h at 65°C in Elusion buffer (20 mM Tris–HCl, pH 7.5, 5 mM EDTA, 50 mM NaCl, 1% SDS, 100 µg RNase A, and 50 µg proteinase K), and finally cleaned up using Zymo ChIP DNA Clean & Concentrator (Zymo Research). Libraries were prepared using NEXTflex DNA barcodes (Bioo Scientific). ChIP-seq libraries were subjected to 75 cycles of single-end sequencing on the NextSeq 500 system. The data was deposited in the GEO database (GSE279957).

##### ChIP-seq data analysis

FASTQ files of the H3K4me3 histone ChIP-seq data from the 230-238, Scid.adh.2C and *Wt*/*Ebf1-/-* pro-B cells were trimmed with Trim Galore (v.0.6.7) using the following options: -fastqc –clip_R1 5 –three_prime_clip_R1 3. Trimmed reads were mapped to the mm10 reference genome (GRCm38) using Bowtie (v.2.4.4). Bigwig files for genome browser visualization were generated with bamCoverage (–normalizeUsing RPGC –centerReads) from deepTools (v1.9). Peak calling was performed with a custom pipeline (https://github.com/Tingvall/macs2_idr), which included peak calling with MACS2 (v2.2.7.1) followed by IDR (v.2.0.4.2). The pipeline was run in narrow mode and the following options were used: –macs_q 0.05 –idr_threshold 0.05. The IDR optional peak set was used for the downstream analysis.

#### HiChIP and PLAC-sequencing and data analysis

##### PLAC-sequencing

PLAC-seq was carried out in duplicates as previously reported (47). Scid.adh.2C cells were cross-linked with 1% formaldehyde for 5 min, re-suspended in ice cold lysis buffer (10 mM Tris-HCl (pH 7.5) 10mM NaCl, 0.2% NP-40) + Protease Inhibitors Cocktail (PIC, Roche) and incubated 15 min on ice. Pelleted cells were washed in ice-cold lysis buffer + PIC and resuspended in 0.5% SDS and incubated at 62°C for10 min. The reaction was diluted with water and 10% Triton X-100, followed by incubation at 37°C for 15 min. Restriction enzyme digestion with 40U Mbol and 25µl NEB2 buffer was performed by incubation at 37°C for 2h (shaking, 900 rpm), followed by inactivation by 20 min incubation at 62°C. Fill-in reaction was conducted with 0.3 mM Biotin-14-dATP (ThermoFisher, 19524016), 0.3 mM dCTP, 0.3 mM dTTP, 0.3mM dGTP and 40U Klenow (NEB, M0210) at 37°C for 1.5h with shaking, followed by ligation reaction by incubation at RT for 2 h with rotation in ligation mastermix (T4 ligation buffer (NEB, B0202), 1% Triton-X 100, 120 ug BSA (NEB, B9000), 4000U T4 DNA ligase (NEB, M0202)). Pellet cells were resuspended in 250µl ChIP SDS-lysis buffer (0.5% SDS, 10 mM EDTA, 0.5 mM EGTA,50 mM Tris-HCl (pH 8.0)) + PIC followed by sonication on Covaris ME220 for 6 min (Peakpower = 75, cycles per burst = 1000, Duty Factor = 15%). Chromatin was spun spun down at 13000 rpm for 10 min and supernatants were diluted in 750µl HBSS + 1 ml RIPA (20 mM Tris–HCl, pH 7.5, 2 mM EDTA, 2% Triton X-100,0.1% SDS, 0.2% sodium deoxycholate, 200 mM NaCl) + PIC. 1% input was removed and 10 µg H3K4me3 (Millipore, 07-473) pre-absorbed to 60 µl Protein-G dynabeads was added to remining sample, followed by ON incubation at 4°C with rotation. Samples were washed 2x in Low Salt ImmuneComplex Wash Buffer (0.1% SDS, 1 % Triton X-100, 2 mM EDTA, 50 mM Tris-HCl pH8,150 mM NaCl), 2x in High Salt Immune Complex Wash Buffer (0.1% SDS, 1 %Triton X 100, 2 mM EDTA, 50 mM Tris-HCl pH 8.0, 500 mM NaCl), 1x in LiCl Immune Complex Wash Buffer (0.25 M LiCl, 1% Igepal-CA630, 1% sodium deoxycholate, 1mM EDTA, 10 mM Tris-HCl pH 8.0), 2x in TE buffer (10 mM Tris–HCl, pH 8.0,10 mM EDTA) followed by elution of chromatin with two rounds of 100µl elution buffer (1%SDS, 100 mM NaHCO3) with shaking (1500 rmp). Eluded chromatin complexes were reverse cross-linked ON at 65°C with the addition of 250 mM NaCl, 100 µg RNase A (ThermoFisher, EN0531) and 50 µg proteinase K (ThermoFisher, AM2546) followed by clean up using Zymo Research ChIP DNA Clean & Concentrator (Zymo Research). 25 µL of Streptavidin T1 beads (Thermo Fisher,65601) washed in Tween Wash Buffer (5 mM Tris-HCl pH 8.0, 0.5 mM EDTA, 1 M NaCl, 0.05% Tween-20) then resuspended in 50 µL of Biotin Binding Buffer (10 mM Tris-13HCl pH 7.5, 1 mM EDTA, 2M NaCl), were added to samples followed by incubation at RT for 15 minutes with shaking. Beads were recovered using a magnet, washed 2x in Tween Wash buffer and incubated at 55°C for 2min with shaking followed by another wash in T4DNA ligation buffer. Captured beads were resuspended in 100 µl end-repair mastermix (0.5 mM dNTPs (VWR, E636-40UMOLE), 12U T4 DNA Polymerase (NEB, M0203), 50U T4 Polynucleotide Kinase (NEB, M0201), 5U Klenow (NEB, M0210), NEBT4 DNA ligase buffer) followed by incubation at RT for 30 min with shaking. Samples were washed 3x in Tween Wash buffer, incubated at 55°C for 2 min with shaking and washed 1x in NEB2 buffer. Beads were captured, and incubated with100 µl A-tailing reaction mix (0.5 mM dATP (ThermoFisher), 25U Klenow Exo- (NEB, M0212), NEB2 buffer) with shaking, washed 2x with Tween Wash buffer and incubation at 55°C for 2 min with shaking. Beads were washed 1x in 100 µl Fast-Link ligation buffer (Epicenter, LK0750H) followed by NEXTflex DNA barcode ligation (BIOO scientific) using the Fast-link ligation kit (Epicenter,LK0750H) and 3x wash in Tween wash buffer, incubation at 55°C FOR 2 min and finally wash in 100 µl 10mMTris-HCl (pH 8.0). PCR amplification of libraries was performed according to optimal number of cycles determined by qPCR, and beads were collected on magnet. Libraries (supernatants) clean-up was performed with Ampure XP (0.8x sample volume). PLAC-seq libraries were subjected to 2 x 38 cycles of paired-end sequencing on the NextSeq 500 system. The data is deposited as in the GEO database (GSE279961).

##### Preprocessing of PLAC-seq data

H3K4me3 PLAC-seq data generated from *Wt* and *Ebf1-/-* fetal liver pro-B cells were generated and preprocessed according to (13). For the preprocessing of H3K4me3 PLAC-seq data from the Scid.adh.2C pro-T cells, FASTQ files from two replicates were merged and trimmed with Trim Galore (v0.6.7) using the following options: (–paired –fastqc –clip_R1 10 –clip_R2 10 –three_prime_clip_R1 3 – three_prime_clip_R2 3. Trimmed reads were processed through the HiC-Pro pipeline (v2.11.4) (48) for the generation of valid interaction pairs. The HiC-pro processing included alignment to the mm10 reference genome (GRCm38) using Bowtie2 (v2.4.2) (49, 50) and the assignment of mapped reads to an mm10 MboI restriction map.

##### Interaction calling

For the H3K27ac HiChIP data obtained from the K562 cells and for the H3K4me3 PLAC-seq data from the Scid.adh.2C cells, interaction calling was performed with FitHiChIP (v9.1) (51) using the merged validParirs output from the HiC-Pro pipeline with the following parameters: IntType=3, BINSIZE=5000, LowDistThr=10000, UppDistThr= 3000000, UseP2PBackgrnd=0, BiasType=1, MergeInt=1, QVALUE=0.05. The corresponding H3K27ac ChIP-seq peak in broadPeak format (GSM733656) was used as the reference peak file. The merged close PEAK-to-ALL interactions, passing the significance threshold (q<0.05), was used as the input for peak annotation conducted using ICE-A. The H3K27ac ChIP-seq peaks (GSM733656) and H3K4me3 ChIPseq peaks (this study) from matching cell types were provided as PeakFile references. For the Wt and Ebf1-/- cells, ALL-to-ALL interactions were generated with the same parameters, except for IntType=4. For comparison, ALL-to-ALL interactions were also generated for the Scid.adh.2C cells.

##### ATAC-seq data analysis

ATAC-seq data were preprocessed with the ENCODE ATAC-seq pipeline (https://github.com/ENCODEDCC/atac-seq-pipeline) (v.2.2.1). For paired-end data from 230-238 cells, the pipeline was run with the following argument: (“atac.paired_end”:true, “atac.auto_detect_adapter”: true, “atac.multimapping”: 4, “atac.mapq_thresh”: 30, “atac.dup_marker”: “picard”,”atac.no_dup_removal”: false, “atac.enable_idr”: true, “atac.idr_thresh”: 0.05, “atac.enable_count_signal _track”: true, “atac.filter_chrs”: [“chrM”, “MT”]. Single-end data from Scid.adh.2C and *Wt*/*Ebf1-/-* pro-B cells were analyzed as for the 230-238 cells with the following differences: “atac.paired_end”:true “atac.no_dup_removal”: false. The IDR optimal peak set was used for the downstream analysis.

##### RNA-seq data analysis

RNA-seq data derived from common progenitor populations, as well as from early B- and T-cell progenitors from the Immgen consortium (GSE122597) were preprocessed by trimming with TrimGalore (v0.6.4), followed by alignment to the mm10 mouse reference genome (GRCm38.p5.vM15) using STAR (v2.7.3). Transcript levels were generated with rsem-calculate-expression in RSEM (v1.3.0), with the following options: –paired-end –alignments –forward-prob 0 –seed-length 20 –outputgenome-bam –sampling-for-bam –estimate-rspd –calc-c.

##### Identification of functional K562 enhancers

CRISPRi-FlowFISH-validated enhancer-promoter pairs in K562 cells (52) were used to assess the ability of ICE-A to annotate cis-regulatory elements compared to other available annotation tools [proximity annotation with HOMER (43) and GREAT (37). Enhancer-promoter pairs [taken from Supplementary Table 3a in the paper of Fulco et al. (2019)] (52) were filtered to exclude: (i) candidate enhancer elements with a distance >3 Mb from the target TSS; and (ii) elements within promoters or gene bodies. Enhancer-promoter pairs with a reported false discovery rate (FDR) <0.05 were considered significant enhancers with respect to their corresponding target gene(s) and were used as inputs for the evaluation of the different annotation methods. Proximity annotation of validated enhancers was performed using annotatePeaks.pl with HOMER (v4.10.0) using the hg19 RefSeq annotations. Annotation with GREAT (v.4.0) was performed using the Basal plus extension mode with default settings for the basal regulatory domain. The maximum distance for annotation of distal regions was increased to 3 Mb, corresponding to the maximum distance used for interaction calling of the HiCHiP data GSM2705043-GSM2705045 (53). For ICE-A, the interaction-based annotation of the validated enhancers was based on H3K27ac HiChIP significant interactions that were called with FitHiChIP. ICE-A was run in Basic mode with the default parameters, with the exceptions of –multiple_anno keep. For all three annotation methods, the numbers and percentages of correctly annotated enhancers were evaluated and presented, both overall and on the individual gene level for different distance ranges. In addition, the performance of ICE-A annotation without proximal annotation were evaluated and compared against other annotation methods, with regards to the ability to identify significant enhancers and their precision (percentage of functional enhancers among total annotated elements). Fraction of enhancers with H3K27ac was evaluated and WashU Epigenome Browser (v54.0.6) was used to visualize identified enhancers of *Gata1*, along with the H3K27ac ChIPseq and HiChIP data.

##### Annotation of enhancers for cancer fitness

Benchmarking of ICE-A against alternative annotation methods were performed with enhancers essential for cancer fitness and proliferation from Chen et al.(54), obtained from a CRISPRi screening. Essential enhancer in HCT-116 colorectal cancer cell line were filtered for having a TSS of an essential target gene within 2Mb. Target genes discovery of essential enhancers was performed with proximity annotation using HOMER (v4.10.0), GREAT (v.4.0) with default setting and ICE-A (default settings except – close_peak_type distance –close_promoter_type distance and –multiple_anno keep). Hg38 TSS location from GREAT (v.4.0) were used for all annotation methods (specified with -gtf in HOMER and –tss in ICE-A). For ICE-A, H3K4me3 PLAC-seq interactions from HCT-116 cell line, as well as A549 lung cancer cell line was used for annotation (the interactions down sampled to have equal number of total interactions, based on interaction score). In addition, ICE-A with proximity annotations excluded were compared against the other annotation strategies. The performance to identify relevant target genes were evaluated based on fraction of essential target genes assigned to at least one essential gene. In addition, the specificity was evaluated based on fraction of essential genes among all target gene assignments.

##### Investigation of lineage-specific elements

###### B- and T-lineage specific elements

A gene-level matrix from the RSEM expression tool was used to estimate the numbers of progenitor and lymphoid populations from the Immgen consortium (LTHSC_CD34-, LTHSC_CD34+, STHSC, MPP4, CLP, FrA, FrBC, FrE, DN1, DN2a, DN2b, DN3) using tximport (v1.3.0), and then variance stabilizing transformed using DESeq2 (v1.42.0). K-mean clustering and visualization of z-scored variance stabilizing transformation (VST) counts averaged per population were performed using the ComplexHeatmap package (v2.18.0). The clustering was restricted to genes that were identified as being differentially expressed between committed FrBC B-progenitor and DN2b T-progenitor cell populations (padj <0.01 & |log2FC|>2) using DESeq2. Clusters with a lineage-specific expression profile were used to define B- and T-lineage-specific gene sets. Lineage-specific elements were defined as open regions from the 230-238 B-cell progenitor cell line (GSE162858) or Scid.adh.2C2 T-cell progenitor cell line (GSE93755), which interacted with the corresponding lineage-specific gene sets. Assignment of target genes to open chromatin regions was based on matching H3K4me3 PLAC-seq interactions using ICE-A with the default options, with the exceptions of: –close_peak_type distance –close_promoter_type distance. Distal elements are defined as non-promoter regions.

###### Chromatin accessibility and motif enrichment

ATAC-seq counts in open chromatin regions for 230-238 B-cell progenitor cell line (GSE162858) and Scid.adh.2C2 T-cell progenitor cell line (GSE93755) were extracted and normalized based on full library size using DiffBind (v3.12.0). Replicates are aggregated and log2 mean normalized counts are visualized in a violin plot for each set of lineage-specific distal elements using ggplot2 (v3.5.1). Statistical analysis is performed with stat_comapre_means from the ggpubr package (v0.6.0) and based on Mann–Whitney U test. The difference in chromatin accessibility was also evaluated separately based on annotation strategy used for target gene assignment. De novo motif enrichment analysis is performed for B-cell- and T-cell-specific distal elements using find-MotifGenome -size 200 -mask from HOMER (v4.10.0). The chromatin accessibility in B- and T cell associated elements were also evaluated based on type of annotation (proximity and/or interaction-based annotation) used by ICE-A.

###### Transcription factor co-occupancy analysis

Co-occupancy of the lymphoid TFs EBF1 (GSE159957), PAX5 (GSE126375), TCF7 (GSE131673) and GATA3 (GSE93755) in lineage-specific elements is investigated using ICE-A run in multiple mode (–mode multiple) with the following options: –upset_plot –circos_plot – circos_use_promoters –skip_promoter_promoter. The co-occupancy analysis is restricted to lineage-specific distal elements (–in_regions regions.bed), interacting with B- and T-cell gene sets (–filter_genes –genes genes.txt).

#### Regulation of broadly expressed genes

##### Investigation of Ebf1 target genes

EBF1 target genes are defined as genes associated to EBF1 bound elements with ICE-A annotation (default options used, with the exceptions of: –close_peak_type distance – close_promoter_type distance). The gene expression patterns of EBF1 targets in early lymphoid development were visualized in a heatmap of the Immgen gene expression data, as described for the identification of B-/T-cell-specific elements, except that EBF1-associated genes were selected. EBF1 targets were classified based on differential gene expression between FL Wt and Ebf1-deficient pro-B cells from GSE92434. Normalized gene expression data were merged per cell type and filtered for genes with normalized score >1 in at least one of the cell types. Up-regulated or down-regulated genes in Wt vs Ebf1-/- were defined as genes with a significant difference in gene expression (padj <0.05) and |log2FC| >1, and common genes were defined as non-differentially expressed genes with |log2FC| <log2(1.5). CompareClusters from the clusterProfiler package (v4.10.0) was used to compare the levels of enrichment of biological process gene ontologies between the different categories of EBf1 targets, split into genes that are bound by EBF1 in promoter or distal (>2.5 kb from TSS) elements.

##### Ebf1-mediated chromatin accessibility change

ATAC-seq data from *Wt* and *Ebf1* -deficient FL pro-B cells (GSE92434) were used to investigate the effect of EBF1 activity on chromatin accessibility. A consensus peak set for the Wt and Ebf1-/- data, defined as the aggregated union of peaks, was generated using bedtools (v2.30.0). ATAC-seq counts for the *Wt* and *Ebf1-/-* pro-B cells in the consensus peak set were extracted using DiffBind (v3.12.0). Differential accessibility analysis between Wt and Ebf1-/- cells was performed using DESeq2 (v1.42.0). The percentage of elements with a significant EBF1-mediated change in chromatin accessibility (|log2FC|>1, padj <0.05) were visualized in a stacked barplot. The analysis was restricted to elements that were open and associated with at least one gene in Wt pro-B cells, split into different categories based on EBF1-mediated changes in gene expression and type of element (promoter/distal). Comparative motif analysis of elements split into categories of EBF1 occupancy and direction of change was performed with gimme maelstrom from GimmeMotifs (v0.15.2).

##### EBF1-addicted gene regulation

To investigate a potential scenario of EBF1-addicted gene regulation, gene expression data derived from the EBF1 degradation system of Zolotarev et al. (55) were used. Genes with EBF1-addicted activation or repression were defined as commonly expressed genes in Wt vs Ebf1-/- pro-B cells, with significantly differential expression at EBF1 degradation (padj <0.05, |FC| >1.5). Gene ontology enrichment analysis of the biologic processes for the top 50 genes (based on log2FC in the degradation system) with EBF1-addicted activation/repression was performed using CompareClusters from the clusterProfiler package (v4.10.0). Myc represent an example of a gene with EBF1-addicted activation. The WashU Epigenome Browser (v54.0.6) was used to visualize Ebf1 binding, chromatin accessibility of *Wt* and *Ebf1-/-* pro-B cells, and H3K4me3 PLAC-seq interactions from Wt pro-B cells at the Myc locus, with a zoomed-in view of the BENC region located 1.6 Mb upstream of the TSS.

#### Inactivation of putative EBF1 target genes

##### Virus expressed guides for CRISPR/Cas9

CRISPR guides were identified by overlaying the genomic coordinates of coding genes of interest with potential CRISPR/Cas9 guides in Benchling (Biology Software. 2020) using the single guide, mm10 genome and 3’ NGG PAM settings. Guides which promote Cas9 mediated cutting within the coding regions were selected for cloning into pAW13.lentiGuide-emCherry plasmid (lentiguide-mCherry, Addgene plasmid 104375; http://n2t.net/addgene:104375; RRID: Addgene_104375, a gift from Richard Young). Lentiviruses were produced by transfecting 293T-HEK cells with lentiguidemCherry plasmids as well as psPax2 and pMD2G packaging plasmids together with X-tremeGENE HP DNA Transfection Reagent (Sigma) according to the manufacturer’s instructions. The resulting virus was harvested after 54-64 hours and concentrated using the Lenti-X concentrator according to the manufacturer’s instructions (Takara Bio).

##### Functional screen for target gene relevance

Hind bones from iCas9 mice aged 8-16 weeks were crushed in a mortar and the cell suspension passed through a 50um filter. KIT+ cells and subsequently purified by magnet-activated cell sorting column using anti-CD117 immunomagnetic beads (Miltenyi Biotec). The cells were transduced with lentiviruses carrying gRNAs targeting putative EBF1-target genes as well as positive (Ebf1) and negative (Rosa 26, R26) control gRNAs. Briefly, non-tissue coated plates were coated with 40 µg/ml retronectin (Takara Bio) overnight at 4°C, blocked with a 2% BSA solution at RT for 30 minutes and washed with PBS. Thereafter, viruses were added, and plates spun at 2000 x g and 32°C for 2 hours. The remaining viral supernatant was aspirated, and wells were washed with PBS. KIT+ cells were then added to the virus-coated wells and plates spun at 300 x g and 32°C for five minutes. After an overnight culture in OptiMEM media supplemented with 10% fetal calf serum, 50µg/mL gentamicin, 50µM β-mercaptoethanol, 50ng/mL KIT ligand, 50ng/mL Fms-like tyrosine kinase 3 ligand (FLT3L) and 50ng/mL interleukin-7 (IL-7) and Doxycyclin (DOX, Sigma) to induce Cas9 expression, transduced BM cells were put in co-culture with OP9 or OP9D stromal cells. The cells were cultured in OptiMEM media supplemented with 10% fetal calf serum, 50µg/mL gentamicin, 50µM β-mercaptoethanol, 10ng/mL KIT ligand, 10ng/mL FLT3L and 10ng/mL IL-7 and DOX (0.1 µg/mL DOX every two days). At day3 post transduction, CD45+mCherry+ cells were sorted for subsequent OP9/OP9D co-culture as above. At day 11, the cells were stained with an antibody cocktail containing CD45 (30-F11, AF700, BioLegend, 103128), CD19 (1D3, PECy7, eBioscience, 15360900), Thy1.2 (53-2.1, Pacific Blue, BioLegend, 140306), Gr1 (RB6-8C5, APCCy7, BioLegend, 108424), CD11b (M1/70, APCCy7, BioLegend, 101226), NK1.1 (PK136, PE, BioLegned, 108708) and CD11c (N418, APC, BioLegned, 117310). 7AAD was used as a viability marker. The absolute number of B-cells (CD45+Gr1-CD11b-CD11c-NK1.1-CD19+) and T-cells (CD45+Gr1-CD11b-CD11c-NK1.1-CD19-Thy1+) generated was determined by FACS analysis using the addition of a set amount of CountBright Absolute Counting Beads (Fisher Scientific) and presented relative to R26 control.

#### Dynamics of promoter/enhancer interactions

##### Interaction landscape in lymphoid development

A consensus set of distal elements involved in lymphoid development (based on open chromatin regions in 230-230 or Scid.adh.2C2 cells; see Figure 2) were annotated in a lineage-specific manner using ICE-A with H3K4me3 significant PLAC-seq interactions from Scid.adh.2C2 pro-T cells and Wt/Ebf1-/- pro-B cells. For comparison, the numbers of total ALL-to-ALL interactions were subsetted to the dataset with the fewest interactions (top interactions based on FitHiChIP q-values are retained). Distal elements were filtered for interactions with B- or T-lineage-specific genes (defined in Figure 2) in any of the cell types. For these putative lineage-specific DREs, the overall enhancer-promoter interaction landscape was compared between cell types and visualized with Circos plots using the circlize package (v0.4.16). Only interactions with B-/T-lineage-restricted genes (defined in Figure 2A) or common target genes [|log2FC FrBCvsDN2b | <log2(1.5)] with activated promoters were included (defined by overlap with H3K4me3 in the promoter of the respective cell type). For visualization of the dynamics of enhancer/promoter interactions in early lymphocytic development, a Venn diagram displaying the numbers and percentages of interactions with B- or T-lineage-restricted genes present in each cell type was generated using the ggVennDiagram package (v1.5.2). In addition, a Venn diagram displaying the degree of overlap between the interactions with alternative common target genes for the lineage-restricted, gene-associated DREs was generated.

##### Genome browser visualization

Genome browser tracks of interactions associated with putative lineage-specific DREs were generated for visualization in the WashU Epigenome Browser (v54.0.6), by compressing and indexing sorted and filtered H3K4me3 PLAC-seq interactions from the Scid.adh.2C2 and FL *Wt* and *Ebf1-/-* pro-B cells using bgzip and tabix from htslib (v1.10.2). The filtered interactions, along with ATAC-seq and H3K4me3 tracks from each cell type, as well as EBF1 and PAX5 binding were visualized in the WashU Epigenome Browser (v54.0.6) at the *Gata3* locus.

### QUANTIFICATION AND STATISTICAL ANALYSIS

Analysis workflows for HiChIP/PLAC-seq, ChIP-seq, ATAC-seq and RNA-seq are presented in detail in the method details section. For chromatin interaction data, FitHiChiP (v9.1) is used for calling significant (q-value<0.05) interactions. For ChIP-seq/ATAC-seq data peaks are called using macs2 (v2.2.71) with a q-value threshold of 0.05. Significant peaks are defined as peaks passing an irreproducibility discovery rate (IDR) of 0.05 from two replicates. For ChIP-seq matching input controls are used in peak calling. Transcript levels count from RNA-seq data were generated with rsemcalculate-expression in STAR (v1.3.0). Differential accessibility and expression analysis is performed with Wald test using DESeq2 (v1.42.0). Features with an Benjamini-Hochberg adjusted p-value < 0.05 are considered significant is not specified otherwise. Log2FC cutoff value for specific analysis are provided in the respective figure legends. Motif enrichment analysis is performed with HOMER (v4.10.0) for de novo motif analysis or GimmeMotifs (v0.15.2) for comparative motif enrichment. Gene set enrichment analysis was performed with the CompareCluster from the clusterProfiler package (v4.10.0) with a significance level of padj (Benjamini-Hochberg correction) < 0.05. If not stated otherwise, statistical analysis is performed with the stat_comapre_means from the ggpubr package (v0.6.0) and based on Mann–Whitney U test. Statistical significance levels are denoted as follows, p-value <0.05: *, p-value <0.01: p**, p-value<0.001: ***, p-value<0.0001: ****.

### ADDITIONAL RESOURCES

ICE-A is available as an open source tool on GitHub: https://github.com/Tingvall/ICE_A.

## Results

### Interaction-based annotation allows for cell type-specific identification of cis-regulatory elements

Current standards for the annotation of DREs to target genes are often based on the concept of proximity, defined as the distance between the DRE and the target promoter in the linear DNA (Figure S1D). Thus, the ability of the conventional annotation of DREs, being based on proximity and GREAT, is dependent upon the local gene density and distances to neighboring genes. To determine how this affects the possibility to identify distal elements, we calculated the theoretical upper distance limit for assigning distal elements to each gene in the human (hg38) and mouse (mm10) genomes. Based on the method-specific annotation rules (i.e., half the distance to a neighboring gene for the standard proximity annotation and the default basal plus extension rule in GREAT), we determined the median distance threshold to be 35 kb or 67 kb for the mouse genome and 47 kb or 91 kb for the human genome, for the standard proximity annotation and GREAT, respectively. If directionality is considered, the ability to identify distal gene-regulatory elements is even more-restricted in terms of upstream and downstream distances from the TSS (Figure S1A). Even though there are gaps in the knowledge regarding DREs, estimates suggest that the median distance for enhancer-promoter pairs is in the range of 100–500 kb (14, 18–20). Taking the constraints associated with proximity-based annotation into consideration, it follows that using the rather conservative estimate of 100 kb, it is possible to identify DREs for less than one-third of the genes in the human genome when applying the standard proximity-based annotation (Figure S1B). Thus, there is strong motivation to develop more-advanced methods for the assignment of DREs to target genes.

To facilitate the annotation of DREs to their target genes, we developed ICE-A (Figure S1C-D), based on the reproducibility-focused workflow management system Nextflow (40). ICE-A incorporates chromatin interaction data into the annotation process using 2D-bed files, making it compatible with several different interaction-calling software’s. To counteract some of the negative features associated with using predefined interactions for peak annotation, such as bin size and minimum distance for interaction calling, ICE-A allows for combined usage of interaction-based and proximity-peak annotation systems. The main output from ICE-A is a single text file for each provided input bed file with information about each annotated element, including the gene symbol, entrezID, distance to the TSS for each assigned target gene, type of annotation used (proximal or interaction-based), and the interaction score from the 2D-bed file (if provided). To allow for the incorporation of different types of data, ICE-A has three different modes. The Basic mode performs interaction-based annotation of a single bed file. The Multiple mode is suitable in cases where the overlap between sets of regions (e.g., co-occupancy of multiple transcriptional regulators) is of interest for the analysis. In addition to performing annotation of every set of regions, the Multiple mode identifies and visualizes overlaps in the form of an UpSet plot or an interaction-based Circos plot. The third mode, integrates gene expression data, allowing for annotated elements to be associated with changes in gene expression. As ICE-A works with pre-processed data files, run times are short. A run using basic mode with four peak files takes approximately two minutes and in multiple mode, including generation of an upset plot, eight minutes, on an eight-core laptop. Thus, ICE-A is an analysis tool that facilitates the integration of different types of sequence-based ‘omics data, making it highly suitable for the analysis of Gene regulatory elements (GREs).

To compare the performances of ICE-A and proximity-based methods for the basic annotation of DREs to target genes, we used CRISPRi-FlowFISH data from functionally validated enhancer-gene pairs in the myelogenous leukemia cell line K562 (52). In that paper, the functionalities of enhancers based on their involvement in transcription were determined in an unbiased way, by evaluating the functional impacts of all open chromatin regions within 450 kb of the TSS as potential regulatory elements (52). The interaction-based annotation of distal elements with ICE-A were based on identification of significant interactions called from H3K27ac HiChIP data (GSM2705043-45) generated from K562 cells (53). This combination of datasets has previously been used for benchmarking of other tools for chromatin interaction processing and analysis (51, 56) as this histone mark can be found at the majority of the active enhancer elements (Figure S1E). All the evaluated annotation tools identified most of the regulatory elements that were located <10 kb from the TSS (Figure 1A). However, for the annotation of distally located enhancers (>50 kb from the TSS), ICE-A was superior to the two proximity-based annotation methods (Figure 1A). ICE-A identified at least one distal enhancer (>10 kb from the target TSS) for 15 of the 17 genes with experimentally validated distal enhancers, as compared with 5 and 0 genes for the GREAT and HOMER proximity annotation, respectively (Figure 1B). Using ICE-A with inactivated proximity annotation function reveled that about 30% of the elements located within 25 kb from the TSS lost annotation (Figure 1D) resulting in reduced efficiency in DRE annotation (Figure 1E). The effect on annotation of elements located more than 25 kp from the TSS was marginal (Figure 1D, E).

**Figure. 1.**
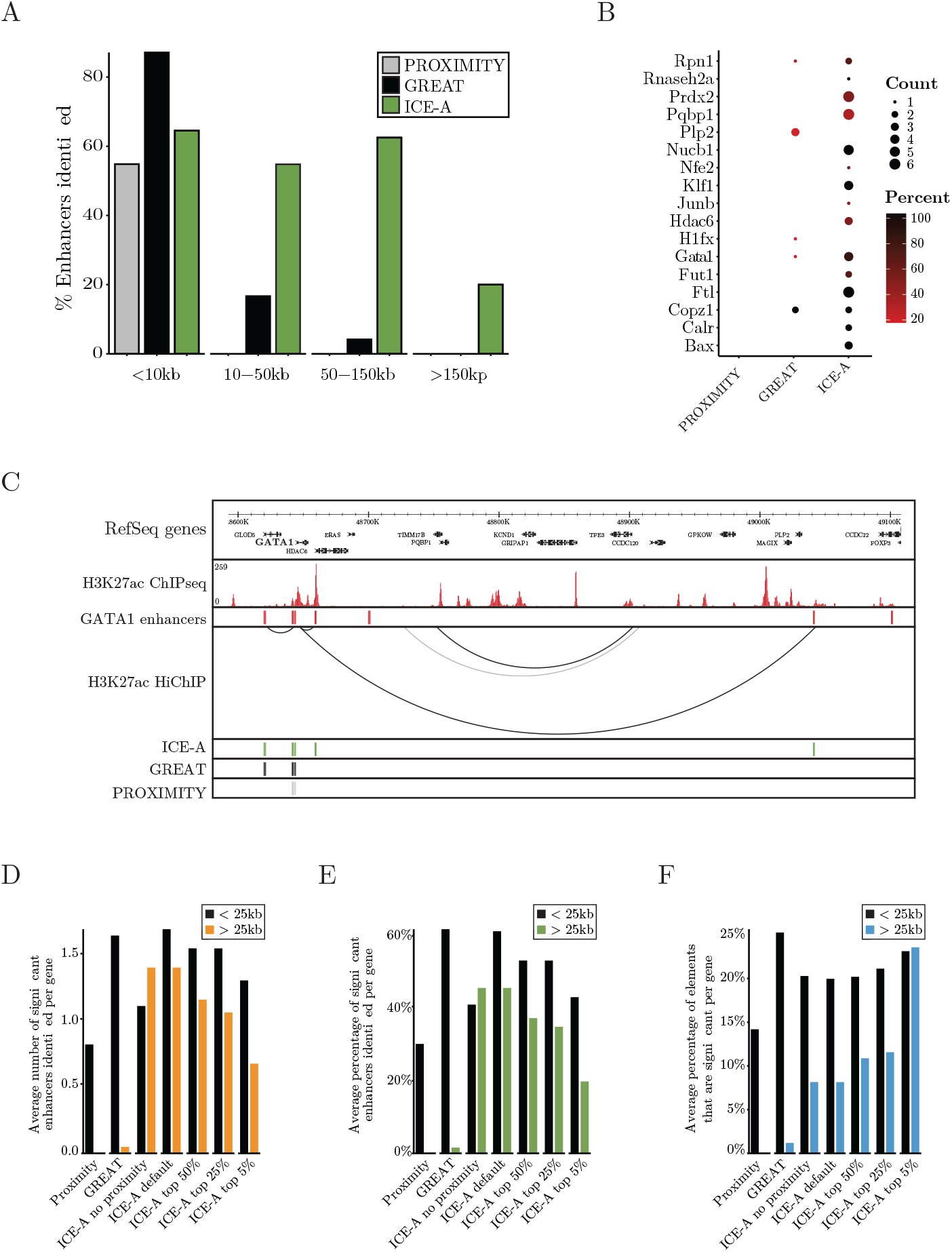
Interaction-based annotation of DREs has advantages over proximity-based approaches. A-C. CRISPRi-FlowFISH -validated enhancers from Fulco et al. (2019) are used for comparisons of the HOMER proximity annotation, GREAT and ICE-A, with respect to their abilities to annotate distal elements. For the ICE-A interaction-based annotation, FitHiChIP significant interactions (q-value 0.05) based on H3K27ac HiChIP (GSM2705043-45) and ChIP-seq (GSM733656) data were used. **A**. Fractions of validated enhancers that were correctly annotated to their corresponding target genes using the different annotation methods. The percentages of identified enhancers are reported for different distance intervals from the relevant TSS. **B**. Comparisons of the numbers and percentages of identified functional distal enhancers (>10 kb from the TSS) per gene, for the HOMER proximity annotation, GREAT and ICE-A. **C**. Example of enhancer annotation at the Gata1 locus. The H3K27ac ChIP-seq, CRISPRi-FlowFISH -validated Gata1 enhancers, and H3K27ac HiChIP (filtered for validated enhancer-promoter interactions) data are visualized in the WashU Epigenome Browser. For each annotation method, the identified enhancers are presented. **D-F**. Average number (D) and percentage (E) of significant enhancers from Fuclo et al. identified per gene from using different annotation methods. F. Average percentage of total elements assigned to each gene with a significant impact on target gene expression. Annotation methods explored in F-H are as follows: Proximity (HOMER), GREAT, ICE-A default, ICE-A no proximity (annotations based on proximity annotation excluded), ICE-A top 50% (top 50% most significant interactions), ICE-A top 25% (top 25% most significant interactions), ICE-A top 5% (top 5% most significant interactions).

The advantage of using ICE-A interaction-based annotation over the proximity-based methods is exemplified by the GATA1 locus (Figure 1C). ICE-A identified three out of the five CRISPRi-FlowFISH validated enhancers, including a distal element located 400 kb downstream of the TSS, as compared to zero and one element for the standard proximity-based annotation and GREAT, respectively. Even though ICE-A was more efficient than proximity-based methods, it could not identify all the functional enhancer in the reference data set (Figure 1A-C). One possibility could be that the H3K27ac based HiChIP analysis did not efficiently detect distal enhancers lacking this histone mark (Figure 1C). To estimate how efficiently H3K27ac marks functional enhancers in the K562 reference data set, we determined the frequency of functional enhancers carrying this histone mark. This revealed that 86% of the functionally relevant elements carried this histone mark (Figure S1E). To estimate the precision in the different methods, we determined how large fractions of the enhancers annotated that was proven functionally relevant in the context of the 562 cells (Figure 1F). Using conventional proximity annotation, about 14% of the distal enhancers located within 25 kb of the TSS, were proven to be functionally relevant. Using GREAT annotation, this frequency was increased to 25% while using ICE-A in a default setting identified 20% of functional elements. Exploring the ability to annotate elements located more than 25 kb from the TSS revealed that only ICE-A could efficiently identify such elements with an efficiency of about 7% in a default setting (Figure 1F). The frequency of functional enhancers annotated could be increased by focusing on the highest ranked elements (Figure 1F). Hence, even if the precision of ICE-A was somewhat lower than for GREAT for identification of functional distal elements located within 25 kb of the TSS, ICE-A was outstanding regarding identification of more distally located elements. Hence, the ability of ICE-A to identify a higher number of functional regulatory elements (Figure 1B), does not come with a dramatic reduction in precision.

To test the efficiency of ICE-As ability to identify functional enhancer elements in a different cellular context, we analyzed an independent dataset assigning regulatory elements to essential genes in cell lines 45 (Figure S1F). ICE-A was able to identify 2-3 times as many functional enhancer gene pairs in the colon cancer cell line (HC1116) as compared to GREAT or proximity annotation methods (Figure S1G). Basing the analysis on Hi-Chip data from a lung carcinoma cell line (A549) or using ICE-A without proximity annotation impaired the ability of ICE-A to identify relevant control elements (Figure S1G). As seen in the analysis of the K562 dataset, the larger number of enhancer/gene pairs identified using ICE-A did not impair the precision of the annotation analysis (Figure S1H). Thus, ICE-A allows for a more-comprehensive annotation of DREs than the current standard methodology.

### ICE-A identifies key regulatory networks in lymphocyte development

The 3D structure of the genome is not static, and chromatin interactions linking a distal enhancer to a target gene can be dependent upon the cellular state or developmental stage (57–60). One advantage of the chromatin interaction-based annotation is the ability to incorporate this dynamic aspect of genome organization into the annotation process, to identify target genes for DREs in a cell type-specific manner. Having developed an efficient tool for interaction-based annotation of DREs, we used lineage-specific target gene assignment of DREs to explore the GRNs active in early lymphocyte development. To this end, we took advantage of the gene expression data for hematopoietic cells generated within the frame of the Immgen consortium, to identify genes that are selectively expressed in B-cell development, as compared to those expressed in T-cell development (Figure 2A, Table S1B). To annotate regulatory elements to these genes in their respective cell types, we used the ICE-A annotation based on our previously published H3K4Me3 PLAC-seq data from the B-cell progenitor cell line 230-238 (GSM4964247). For the annotation of elements to T-lineage genes, we generated H3K4Me3 PLAC-seq data from the T-cell progenitor cell line (Scid.adh.2C2). Exploring the ATAC accessibility of DREs annotated to B-lineage-restricted cells (B-DRE) in B-cell progenitors (pro-B cells), as compared to T-cell progenitors (pro-T cells), revealed generally larger accessibility in B-lymphoid cells (Figure 2B). The opposite was observed for DREs annotated to T-lineage genes (T-DREs). Even though a major part of the elements were identified based on the interaction data (Figure S2A), a set of elements displaying lineage restricted accessibility was identified via the proximity function (Figure S2B). Determining how large fraction of the annotated elements that were identified by proximity as compared to interaction data, supporting the idea that ICE-A can identify lineage-restricted regulatory elements (Figure 2B). To identify the key TFs of the GRNs in B-cell and T-cell progenitors, we performed motif enrichment analysis of elements that were annotated to lineage-restricted genes by ICE-A (Table S1C, D). The B-DREs were enriched for binding sites for BORIS, ETS (SPI-B) and RUNX proteins, as well as for the lineage-restricted TF EBF1 (Figure 2C). T-DREs were similarly predicted to bind the RUNX and ETS proteins as well as the TCF7 and GATA proteins (Figure 2D). To explore further the abilities of these TFs to bind ICE-A-defined regulatory elements, we took advantage of the multiple mode in ICE-A for the identification of TF occupancy at the identified DREs. We included the ChIP-seq data for the TFs EBF1 and PAX5 in pro-B cells and for TCF7 and GATA3 in pro-T cells. Of the B-DREs, 27% were detected as being bound by EBF1, often in combination with PAX5 (Figure 2E). In addition, EBF1 was found to be bound by 8% of the T-DREs in the pro-B cells (Figure 2F). TCF7 binding was detected at approximately 25% of the T-DREs and at 4% of the B-DREs. The observation that EBF1 and PAX5 bind to a substantial fraction of the T-DREs, as well as at promoters of T-lineage-restricted genes (Figure 2F) may reflect the abilities of these B-lineage TFs to repress alternative lineage programs in B-cell progenitors. Our findings are concordant with the essential functions of EBF1 in B-cell development and TCF7 in T-cell differentiation, suggesting that analysis of ICE-A annotations can be used to identify critical components of lineage-restricted GRNs in development.

**Figure. 2.**
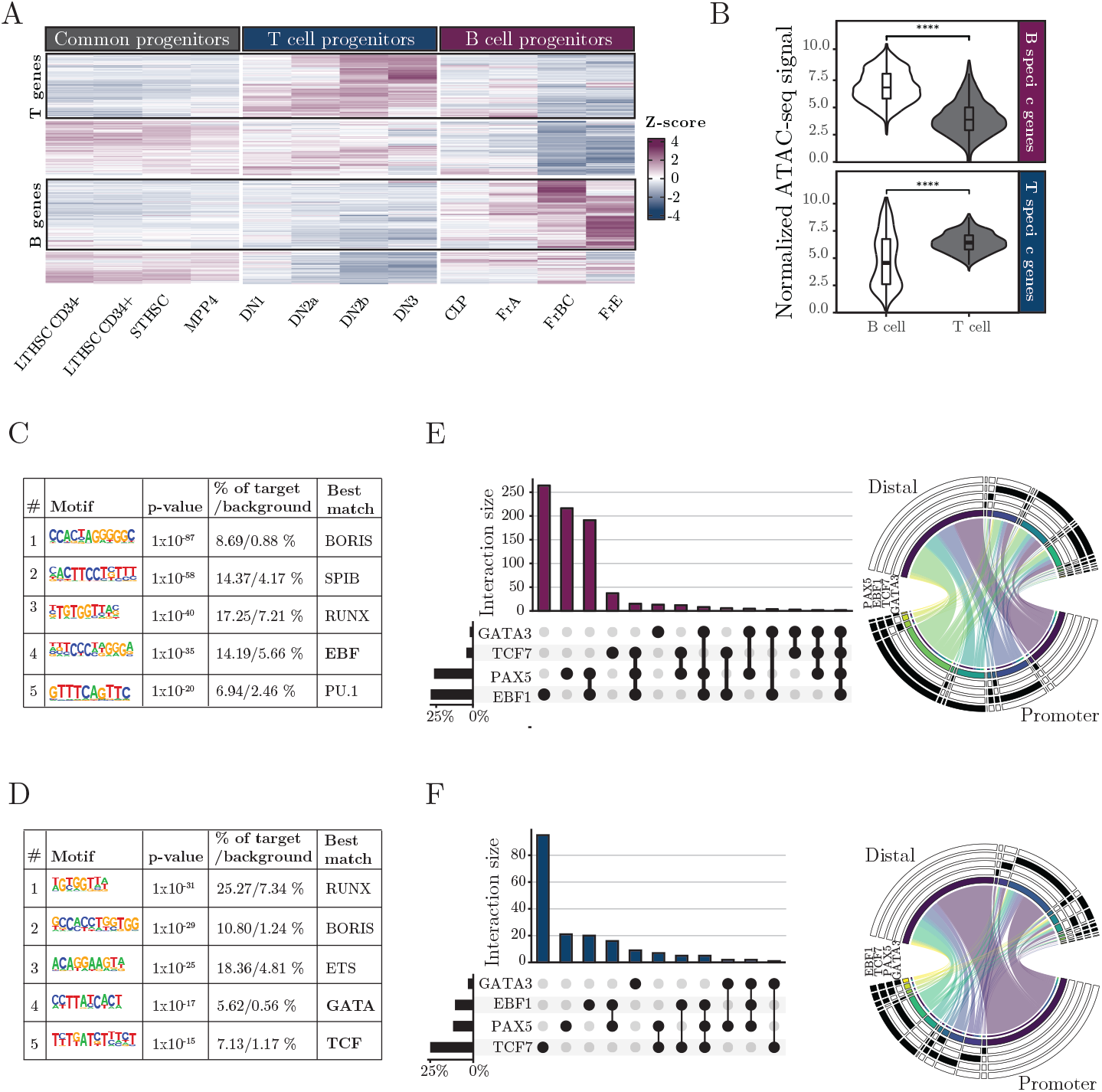
ICE-A interaction-based annotation can identify lineage-specific DREs. **A**. Heatmap with z-scores of the normalized gene expression counts of early lymphoid progenitors from the Immgen consortium (GSE100738). Genes with significantly different expression patterns between FrBC B-cell progenitors and DN2b T-cell progenitors are shown. K-means clustering was used to define the B- and T-cell progenitor-specific gene sets. **B**. Violin plot of normalized ATAC-seq signals of distal elements annotated to a B- or T-lineage gene by ICE-A interaction-based annotation, with H3K4me3 PLAC-seq data from a cell line matching the cell-type-specific gene sets. The ATAC-seq signal levels are compared between B- (230-238) and T- (Scid.adh.2C2) cell progenitor cell lines. Statistical analyses are based on the Mann–Whitney U-test, ****p < 0.0001. **C-D**. Top-rated enriched motifs based on HOMER de novo motif analysis for distal B-(C) and T-(D) cell elements. B- and T-lineage elements are defined as open chromatin regions annotated to B- or T-lineage genes sets using ICE-A with H3K4me3 PLAC-seq interaction in 230-238 and Scid.adh.2C2 cells, respectively. **E-F**. Output from a transcription factor (TF) co-occupancy analysis in ICE-A run in multiple mode, including an UpSet plot of TF co-occupancy of distal elements and a Circos plot of TF overlap in distal and promoter regions of previously defined, cell-type-specific gene sets.

### Lineage-restricted transcription factors interact with the regulatory elements of broadly expressed genes

Even though lineage-specific TFs, such as EBF1 and PAX5, control the expression of lineage-restricted genes, they also appear to be participate in the regulation of genes that are involved in basic biologic processes, such as proliferation, cell survival and metabolism 6-11, (47). To gain a better understanding of how lineage-restricted TFs are integrated into stage and lineage restricted regulatory networks, we used ICE-A to scrutinize GRNs out from a TF-centered perspective, through the identification of target genes annotated to bound regulatory elements. Thus, we identified EBF1-bound promoters and DREs using ChIP-seq data derived from the 230-238 cells, and annotated the bound elements to coding genes using ICE-A. Using gene expression data from the Immgen consortium, we then investigated how these target genes were expressed by hematopoietic progenitors (Figure 3A). While B-lineage genes were identified among the target genes, only 20% (cluster 5) displayed a B-lineage restricted expression pattern and a substantial fraction of the genes were broadly expressed. Performing the same analysis for PAX5, GATA3 and TCF7 yielded highly similar results (Figure S3A) identifying several broadly expressed target genes. The binding of lineage-restricted TFs to regulatory elements annotated to broadly expressed genes suggests that these proteins are integrated into the global GRN of the cell, supporting the idea that these lineage specific TFs control basic cellular processes (6–11, 47).

**Figure 3.**
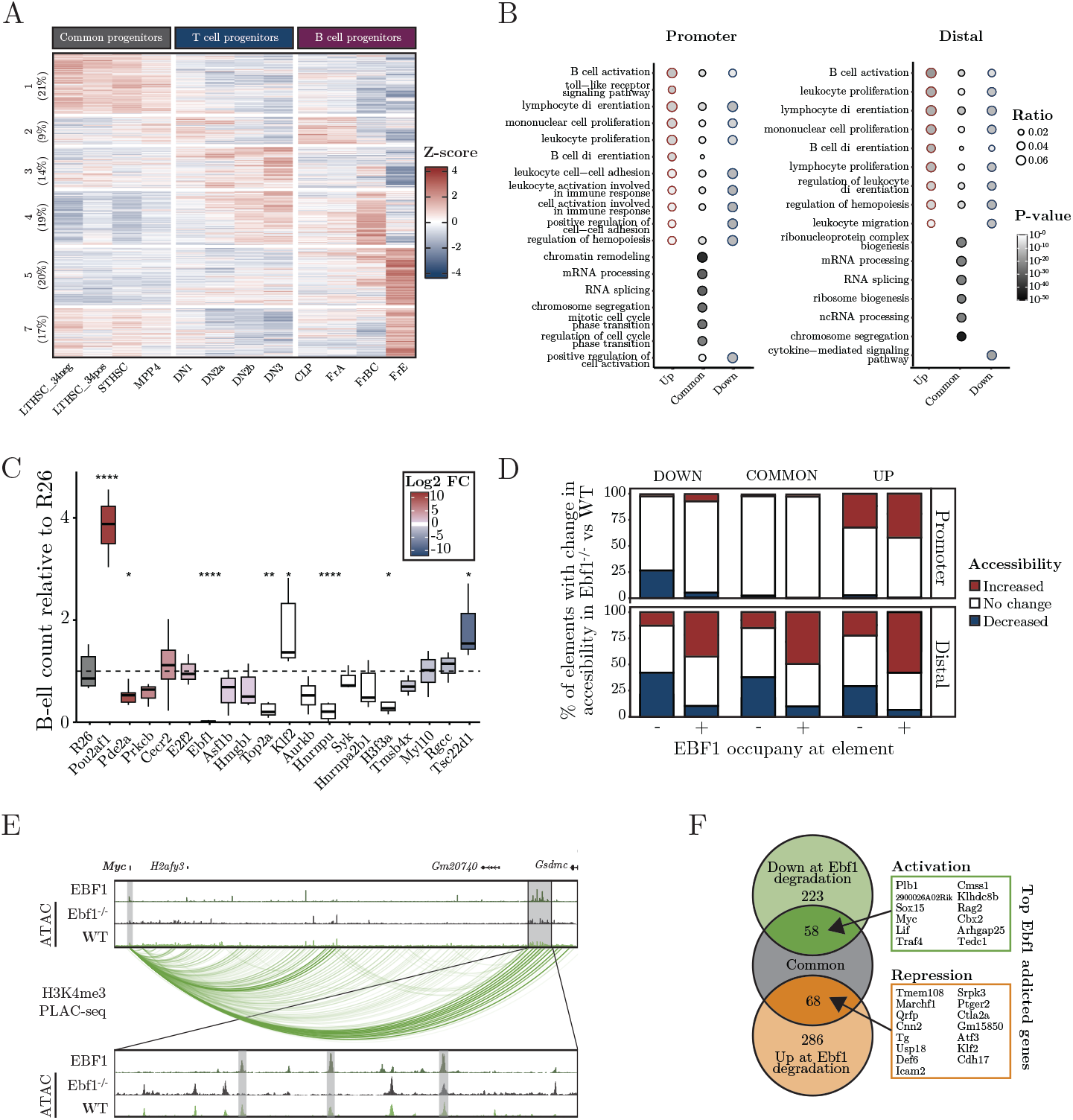
EBF1 interacts with the regulatory elements of broadly expressed genes. **A**. Heatmap with the z-scores of normalized gene expression of Ebf1-bound elements in early lymphoid progenitors from the Immgen dataset (GEO: GSE100738). Percentage of genes in each cluster is presented. **B**. Gene ontology (GO) analysis of enriched biologic processes in genes bound by EBF1 at promoters or distal regions. Gene sets are categorized based on differential expression between Wt and Ebf1-/- FL pro-B cells [Up-/Down-regulated: padj 0.05 & |log2FC| >1, common: log2FC<log2(1.5)]. **C**. Diagram displaying the relative number of generated CD19+ cells after 11 days of in vitro incubation of KIT+ cells on OP9 stroma cells compared to R26 control. The color represents log2FC Wt vs Ebf1-/- from RNA seq data. **D**. Promoter and distal elements associated to the gene sets from 2B are identified by ICE-A interaction-based annotation with H3K4me3 PLAC-seq data from FL Wt pro-B cells. Fraction of elements with a significant change (padj < 0.05 & abs(log2FC) > 1) in accessibility between Wt and Ebf1-/- FL pro-B cells are visualized in a stacked barplot, split into categories based on gene category and EBF1 occupancy. **E**. The Myc locus is shown as an example of a gene that shows EBF1-addicted activation, visualized in the WashU genome Epigenome Browser. Selected EBF1-bound elements located in a distal enhancer region 1.6 Mb from the Myc TSS are highlighted in gray. The epigenome browser tracks include EBF1 ChIP-seq (GEO: GSE159957), ATAC-seq data from Wt/Ebf1-/- FL pro-B cells and H3K4me3 PLAC-seq interactions from Wt FL pro-B cells. **F**. Venn diagram displaying the overlap between common genes (defined as in B) and genes with differential expression at EBF1 degradation (|log2FC| > log2(1.5)), from (GSE201141). Genes that show the strongest EBF1-addicted activation or repressive expression are listed.

To gain further insights into how lineage-restricted TFs, such as EBF1, are integrated into the transcriptional context of a lymphoid progenitor cell, we compared the expression levels of EBF1 target genes in in vitro-expanded *Ebf1-/-* cells and in wildtype Wt pro-B cells. The genes were classified as either up- or down-regulated (padj <0.05 & |log2FC| >1 in *Wt* vs *Ebf1-/-*) or common (|log2FC| < log2(1.5)). Performing a gene ontology analysis, the differentially expressed EBF1 targets, up-regulated or downregulated in *Wt* cells, were enriched for genes linked to GO-terms such as B-cell activation and lymphocyte activation, independently of whether EBF1 was bound at the promoter or at a DRE (Figure 3B). The target genes that were not differentially expressed instead encoded proteins that are involved in basic biologic processes, such as RNA processing and translation (Figure 3B). To determine if broadly expressed target genes are of importance for B-cell development, we used hematopoietic progenitor cells (KIT+ cells) that carry a doxycycline-responsive CAS9-encoding gene. The cells were transduced with lentiviruses carrying a florescent reporter gene (mCherry) and express guides, that target genes annotated to EBF1-bound elements. After sorting for mCherry+ cells, the progenitors were seeded onto OP9 stroma cells in the presence of interleukin (IL)-7, FLT3/FLK-2 ligand (FL) and KIT ligand (KL), and the numbers of CD19+ cells generated in the cultures were determined by flow cytometry. While inactivation of the gene encoding KLF2 resulted in increased generation of CD19+ cells, targeting the histone-encoding *H3f3a*, the ribonuclear proteins *Hnrnpa2b1* and *Hnrnpu* or the DNA topoisomerase *Top2a* genes resulted in reduced generation of B-cell progenitors (Figure 3C). To determine if any of these EBF1 target genes are important for T-cell development, we incubated the progenitor cells on a NOTCH ligand expressing stroma cell, OP9 Delta-1 (OP9D) under conditions allowing for the combined generation of B and T-lineage cells. While the generation of B-lineage cells was impaired by the down-regulation of the same target genes as observed in the OP9 supported B-cells cultures, the generation of T-lineage cells were only significantly impaired upon targeting of *Top2a* (Figure S3B). Therefore, several broadly expressed EBF1 targets genes are of importance for the generation of B-lineage cells.

To attain a better understanding of how EBF1 is integrated into the GRNs of broadly expressed genes, we took advantage of the ATAC-seq data from in vitro-expanded *Ebf1-/-* and *Wt* pro-B cells. All the genes were annotated to regulatory elements with ICE-A, and using the EBF1 ChIP-seq data, we identified the elements that directly bound to EBF1 (EBF1+) (Figure 3D). Exploring the changes in accessibility for the elements annotated to genes expressed at higher levels in Wt cells than in Ebf1-/- pro-B cells (Up), we detected increased accessibility (|log2FC| & >log2(1) at 45%–50% of the elements bound by EBF1. While the promoters linked to commonly expressed genes displayed few changes in accessibility, we detected increased accessibility for approximately half of the EBF1-bound DREs, while elements not bound by EBF1, frequently displayed a loss of accessibility. Thus, EBF1 is integrated into the GREs of commonly expressed genes both by binding to already accessible elements and by modifying the epigenetic landscape at targeted DREs. This is exemplified by the Myc gene, where EBF1 interacts with DREs that display both EBF1-dependent and EBF1-independent accessibility (Figure 3E). This analysis suggests a substantial degree of epigenetic dynamics at elements that are involved in the regulation of broadly expressed genes, and that EBF1-mediated repression may not be accompanied by reductions in epigenetic accessibility at the targeted element.

To investigate the functional impact of EBF1 integration into the GRNs that control the expression of commonly expressed genes, we analyzed data from an experiment in which endogenous EBF1 was replaced by a protein fused with FKBP (F36V) (55), which resulted in rapid degradation of the protein following the addition of the drug TAG13. Analyzing the RNA expression levels, we detected decreased expression (padj < 0.05 & log2FC > log2(1.5)) of 281 genes, 58 of which belong to the commonly expressed genes (log2FC < log2(1.5) in *Wt* vs *Ebf1-/-* pro-B cells), upon degradation of EBF1 (Figure 3F). Using the same criteria, we found 354 genes whose expression levels were upregulated upon loss of EBF1; of these, 68 were classified as commonly expressed genes.

Thus, lineage-specific TFs can modulate the GRNs controlling the expression of broadly expressed genes, to establish tissue-specific control of basic biologic processes.

### ICE-A annotation reveals the dynamics of promoter/enhancer interactions in early lymphocyte development

Just as promoters may collaborate with several DREs (14, 15), it has been suggested that enhancers can target several promoters (21, 22, 36, 61–63). The comprehensive annotation of DREs using ICE-A opens the exciting possibility to explore the complexity of promoter-enhancer pair interactions in different cellular contexts. To this end, using ICE-A, we identified DREs that were annotated to lineage-restricted genes (B-DREs or T-DREs) (Figure 2). We then determined their interactions with target promoters using H3K4me3 PLAC-seq data from in vitro-expanded Wt or Ebf1-/- pro-B cells, as well as pro-T cells (Scid.adh.2C2) (Figure 4A). In the pro-T cells, we detected 1,186 interactions between DREs and promoters of T-lineage-restricted genes (TPs) (Table S1H), as well as 854 interactions between DREs and promoters of B-lineage-restricted genes (BPs) (Table S1H), and 5,324 interactions between DREs and the promoters of commonly expressed genes (UPs). Examination of the B-DRE and T-DRE interactome in Wt pro-B cells revealed a substantial increase in the number of interactions with BPs and a reduction in the number of interactions with TPs, as compared to what we observed in T-lineage cells. However, the majority (66%) of the detected interactions were, just as in the T-lineage cells, with UPs. The interactome of *Ebf1-/-* pro-B cells appeared to have an intermediate profile, with comparable numbers of interactions between the BPs and TPs and the DREs. To investigate further the dynamics of the interactomes for lineage-restricted DREs, we determined the levels of conservation of the detected interactions in the different the cell types (Figure 4B). Among the 1,186 identified interactions with TPs in T-lineage cells, 42% were exclusively detected in the T-lineage cells, whereas 32% were conserved in the *Wt* pro-B cells. Similar numbers were obtained when determining the interactome of the DREs linked to BPs, as 46% of the interactions were unique to the Wt pro-B cells, while 26% of the interactions were shared between the B- and T-lineages. Hence, even if the interactions between B- and T-DREs and UPs are relatively conserved, lineage specific interactions were easily detectable. For the interactions with UPs, the degree of conservation between lineages was higher than for interactions with lineage-restricted genes, with >50% conservation between B-cell and T-cell progenitors for both B-DREs and T-DREs (Figure 4C). The analysis also showed that many of the lineage-restricted interactions were conserved in the *Ebf1-/-* pro-B cells (Figure 4B), confirming that the EBF1-deficient pro-B cells display an interactome that is intermediate to those of the pro-B and pro-T cells. This is exemplified by the *Gata3* gene. The detected promoter-enhancer interactions are almost identical in pro-T cells and *Ebf1-/-* pro-B cells (Figure 4D). Even though the low level of H3K4me3 at the *Gata3* promoter in the Wt-pro-B cells makes it difficult to determine the actual structure of the locus, we detected interaction to an alternative EBF1 and PAX5 bound promoter from one of the distal *Gata3* elements solely in the pro-B cells. These data underline the complexity of GRNs in early lymphocyte development and underpin the importance of EBF1 for the establishment of the epigenetic landscape during B-cell development.

**Figure 4.**
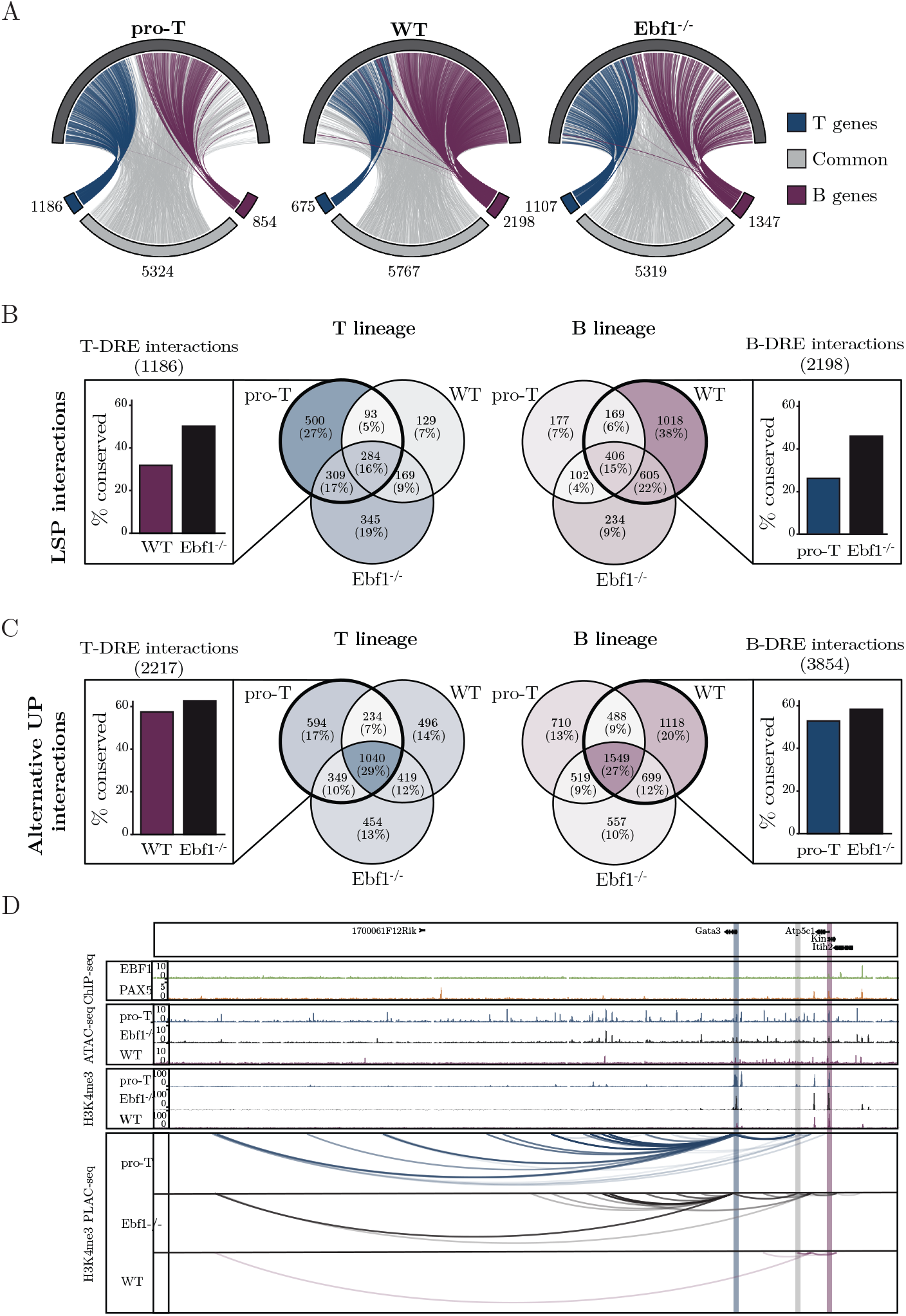
ICE-A annotation reveals the dynamics of promoter-enhancer interactions in early lymphocytic development. **A**. Circos plot of chromatin interactions between putative lineage-specific distal elements (open chromatin regions in B-cell and/or T-cell progenitor cell lines, with associations to the B-cell or T-cell gene list defined in Figure 2A). Chromatin interactions are based on H3K4me3 PLAC-seq interactions for the Scid.adh.2C2 T-cell line, Wt FL pro-B cells and Ebf1-/- FL pro-B cells and are restricted to interactions with H3K4me3-associated promoter regions of common [|log2FC FrBCvsDN2b| < log2(1.5)] genes. Interactions with alternative promoters are included. **B-C**. Venn diagram displaying overlap in DRE interactions to lineage restricted (B) and common (C) promoters. For each lineage, the fraction of conserved interaction from putative lineage-specific distal element (B/T-DREs) in the alternative lineage and Ebf1-/- FL pro-B cells are presented. **D**. Visualization of TF binding, ATAC-seq signal and H3K4me3 status for the Scid.adh.2C2 T-cell line, Wt and Ebf1-/- B-cell progenitors at the Gata3 locus in the WashU Epigenome Browser. H3K4me3 PLAC-seq data from all cell types are included, filtered for interactions associated with putative B- or T-lineage enhancers.

## Discussion

To expand our understanding of stage- and lineage-specific gene activities, there is an increasing need to accurately assign regulatory elements to their target genes. We here present ICE-A, a software that allows for the easy incorporation of chromosomal configuration data to annotate DREs to coding genes. We find that using chromatin interactions to identify putative target genes for distal elements reduces the impact of known limitations that hamper the efficiency of proximity-based methods. In addition, the ability of chromatin capture methods to reflect the dynamic and cell type-specific aspects of genome organization will be reflected in the annotation process and can provide an additional layer of understanding of gene regulatory mechanisms. Even though ICE-A outperforms the proximity-based methods with respect to the annotation of distal enhancers, it is worth noting that GREAT detected the highest number of the most-proximal enhancers (<10 kb from the target TSS) (Figure 1A). This is expected considering the principle of basal gene regulatory domains plus the extension principle applied by GREAT, which allows for the assignment of more than one gene per element. This in contrast to the single closest gene assignment applied in standard proximity annotation as well as in ICE-A for distances below the user-specified interaction threshold. The improved detection rate of GREAT for proximal enhancers, which is linked to the potential for multiple annotations per regulatory region, may come at the cost of an increased rate of “false” annotations. Determining the frequency of elements that could be verified as functionally relevant suggested that about one in five annotated enhancers were important for transcription for the genes in K562 cells. Even though ICE-A may identify DREs that are not critical for the regulation of the gene in question in a given cell type, the annotation is based on direct interactions of the enhancers with the promoter.

However, false positives are inevitable, and the extent to which this poses a significant problem is highly dependent upon the application. ICE-A provides several options that could help tailor the annotation to specific needs, including the option to include only one annotation per region by filtering or ranking of interaction score (Figure 1F), selection or adjustment of the way in which ICE-A handles nearby interactions. Another aspect that gives interaction-based annotation approaches an advantage over proximity annotation strategies is the potential to identify cell type-specific or dynamic regulatory regions. ICE-A does not only outperform existing programs but also allows for easy incorporation of ChIP-seq/Cut&run data, as well as gene expression data in the analytic method allows for complementary approaches that are centered on gene expression patterns, TF binding, the epigenetic landscape or chromosomal interactions. Thus, ICE-A opens new avenues to unraveling GRNs.

Using a gene-centered approach for the identification of DREs annotated to lineage-restricted genes (Figure 2), we were able to identify TFs that act as key regulators of early lymphocyte development (Figure 2). Motif enrichment analysis identified GATA- and TCF7-binding sites in elements annotated to T-lineage genes, while DREs linked to B-cell genes were enriched for EBF1-binding sites. Thus, ICE-A identifies DREs bound by lineage-specific TFs that act as key regulators of early T-cell and B-cell development (1–12). Exploring a TF-centered approach, in which ICE-A annotates elements bound to coding genes, we were able to identify target genes for the TF, thereby providing insights into how specific DNA-bound proteins affect cellular functions. Our analysis of early lymphocytic development suggests that lineage-restricted TFs target regulatory elements that are annotated to many broadly expressed genes. Focusing on the role of EBF1 in B-cell development, several of the bound DREs were annotated to commonly expressed genes, with no significant change in the gene expression levels in *Ebf1-/* - pro-B cells as compared with *Wt* pro-B cells. We provide experimental evidence that several of these EBF1 target genes are of importance for the generation of normal B-cell progenitors, supporting the relevance of these target genes (Figure 3C). There are also several examples of broadly expressed genes that become dependent upon EBF1 once this TF has been expressed (Figure 3D). These genes include Myc (47, 64), which likely explains the significance of EBF1 for the survival and expansion of pro-B cells (10, 47). The mechanism underlying the addiction to EBF1 may involve changes in enhancer or chromatin structure (Figure 3D) (13), or as in the case of the *Myc* gene, activated transcription of a repressor of the target gene (47). Based on our analysis of early lymphocytic development, we believe that ICE-A-based annotation provides unique and relevant insights into GRNs.

The use of ICE-A allows for easy and extensive exploration of the interactome that underlies the general principles of gene regulation in a specific cellular context. Several model systems have provided evidence that DREs can target multiple promoters (21, 22, 36, 61– 63). Our analysis of the DRE interactome in early lymphocyte progenitors suggests that a substantial fraction of the elements that interact with promoters to control the expression of lineage-specific genes may also interact with alternative promoters (Figure 4), often in a cell type-specific manner. While linear proximity appears to account for most of the promoter enhancer specificity in yeast (16) and to 79%–88% of the promoter enhancer specificity in Drosophila (17), only about half of the DREs in a mammalian genome are estimated to follow this principle (21, 22). Therefore, it remains as a persistent challenge to resolve the mechanisms that control enhancer-promoter specificity in complex genomes. In Drosophila, certain DREs show a preference for specific core promoters (65–68) or rely on a proximal “tethering” element that is proximal to the promoter (69–72). Our analysis suggests that lymphocyte progenitors exploit a pool of DREs to control the expression of both lineage-specific and broadly expressed genes, and that the interactome is highly dynamic with enhancers switching to new promoters depending on the cellular context (Figure 5). It has been proposed that alterations in the primary DNA sequence result in the formation of new promoter-enhancer interactions (36, 61, 73–75). Our data indicate that promoter selections may be highly lineage- and stage-specific, possibly because of changes to the epigenetic landscape. Such dynamic changes in DRE-promoter interactions may also contribute to the abilities of TFs to act as both activators and repressors (9, 10, 76–80). This may be of importance in relation to the abilities of lineage-specific proteins such as EBF1, PAX5, GATA3 and TCF7 to interact with both activator and repressor complexes (13, 81), possibly making their functions highly context-dependent. Our analysis of EBF-repressed target genes does, however, suggest that EBF1 binding results in increased accessibility of the DRE, even when it is annotated to a repressed gene. This is concordant with the ability of EBF1 to act as a pioneer factor and to promote the formation of phase-separated structures at target genes (13, 82, 83). While the activation of regulatory elements may appear incongruous with target gene repression, we believe that our findings provide an alternative model for how a TF can contribute to transcriptional repression through diversion of DRE activity from lineage-specific promoters (Figure 5).

**Figure 5.**
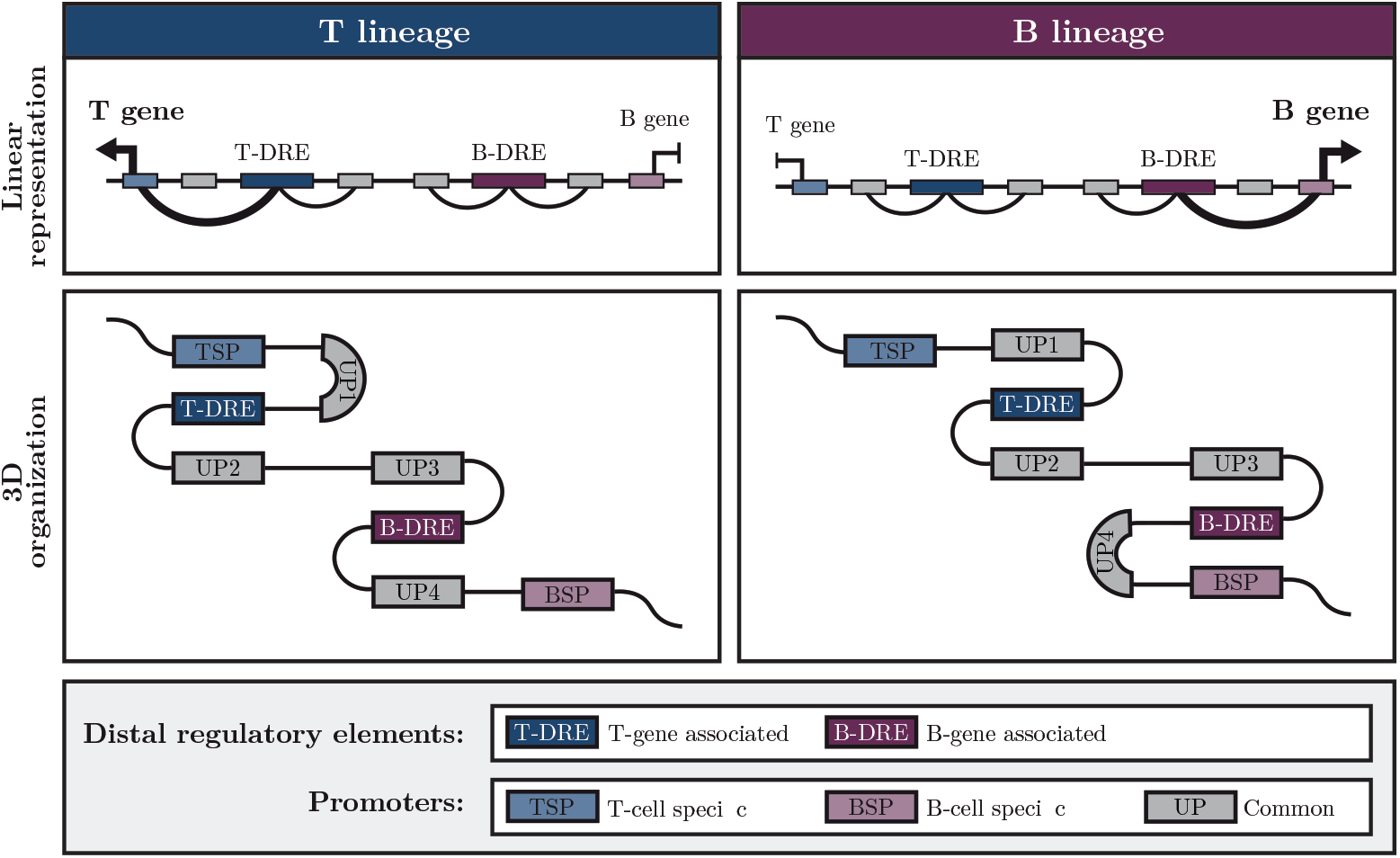
Model of lineage-specific gene regulation in lymphoid development. Schematic representation of a model for lineage specific transcriptional regulation in lymphoid development. Lymphoid progenitors explore a pool of distal regulatory elements (DREs) for control of both lineage specific and common genes. A highly dynamic interactome with distal elements switching to new promoters depending on cellular state and developmental stage, allow for control of both lineage restricted and common genes in a cell type-specific manner.

Our use of ICE-A to dissect the roles of GRNs in early lymphocytic development reveals the versatility of the software and highlights the importance of integrating chromosomal configuration data into the DRE annotation processes. Given that more than 90% of disease- and trait-associated genetic variants reside in non-coding regions of the genome, accurate assignment of DREs to target genes could provide insights into the mechanisms operating in various disease states (84). Therefore, we believe that ICE-A can greatly facilitate the exploration of genomics data in an easy and user-friendly manner.

## RESOURCE AVAILABILITY

### Lead contact

Requests for further information and resources should be directed to and will be fulfilled by the lead contact, Mikael Sigvardsson.

### Materials availability

This study did not generate new unique reagents.

### Data and code availability

- **Data:** The novel sequencing datasets generated for this study are deposited in the Gene Expression Omnibus (GEO) database with accession numbers GSE279957 (ChIP-seq) and GSE279961 (PLAC-seq). The GEO accession number and information related to the previously published data used in this study are presented in Ket Resource Table and Supplementary Table 1A.
- **Code:** ICE-A is available as an open source tool on GitHub: https://github.com/Tingvall/ICE_A. All the code related to the analysis and generation of figures for this article is available on GitHub: https://github.com/Tingvall/ICEA_analysis.
- **Other:** Supplemental information including gene sets, element and guide info is available in Supplementary Table 1.

## ACKNOWLEDGEMENTS

We are grateful for editorial suggestions from Vincent Collins. This work was supported by grants from the Swedish Cancer Society (23-3019P), the Swedish Childhood Cancer Foundation (2022-0019), Stiftelsen för Strategisk Forskning (SSF) (IB23-0001), the Swedish Research Council (2021-02379) and Linköping University.

## AUTHOR CONTRIBUTIONS

JTG in collaboration with JU designed ICE-A. The analysis using ICE-A was performed by JTG in collaboration with MS. The gene targeting KO experiments were performed by CJ. While JTG and MS wrote the first draft of the manuscript, all authors contributed to the finalization of the manuscript.

## DECLARATION OF GENERATIVE AI AND AI-ASSISTED TECHNOLOGIES

AI tools have not been used for the generation of this manuscript

## SUPPLEMENTAL INFORMATION

Document S1: Figure S1-3+Figure legends

Document S2: Excel sheets containing supplementary table S1A-H

