## Supplemental Figures for "Chromatin interaction-based annotation of distal cis-regulatory elements reveals highly dynamic promoter-enhancer interactions in lymphocyte development"

S1

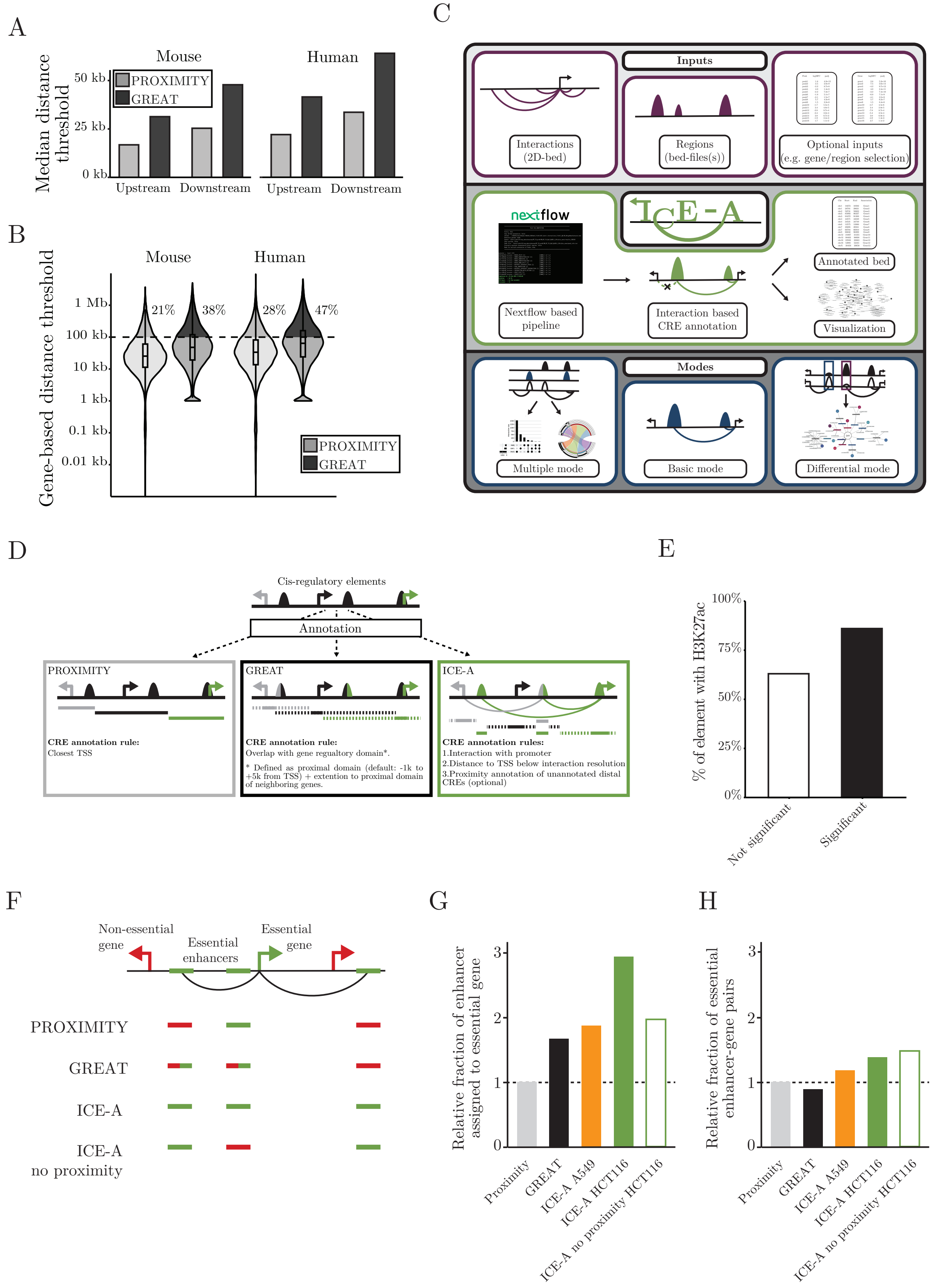
A
Mouse
Human
Median distance threshold
50 kb
25 kb
0 kb
PROXIMITY
GREAT
Upstream
Downstream
Upstream
Downstream
B
Mouse
Human
Gene-based distance threshold
1 Mb
100 kb
10 kb
1 kb
0.1 kb
0.01 kb
21%
38%
28%
47%
PROXIMITY
GREAT
C
Inputs
Interactions (2D-bed)
Regions (bed-files(s))
Optional inputs (e.g. gene/region selection)
nextflow
ICE-A
Annotated bed
Nextflow based pipeline
Interaction based CRE annotation
Visualization
Modes
Multiple mode
Basic mode
Differential mode
D
Cis-regulatory elements
Annotation
PROXIMITY
CRE annotation rule:
Closest TSS
GREAT
CRE annotation rule:
Overlap with gene regualtory domain*.
* Defined as proximal domain (default: -1k to +5k from TSS) + extention to proximal domain of neighboring genes.
ICE-A
CRE annotation rules:
1.Interaction with promoter
2.Distance to TSS below interaction resolution
3.Proximity annotation of unannotated distal CREs (optional)
E
% of element with H3K27ac
100%
75%
50%
25%
0%
Not significant
Significant
F
Non-essential gene
Essential enhancers
Essential gene
PROXIMITY
GREAT
ICE-A
ICE-A no proximity
G
Relative fraction of enhancer assigned to essential gene
3
2
1
0
Proximity
GREAT
ICE-A A549
ICE-A HCT116
ICE-A no proximity HCT116
H
Relative fraction of essential enhancer-gene pairs
3
2
1
0
Proximity
GREAT
ICE-A A549
ICE-A HCT116
ICE-A no proximity HCT116

**Supplementary Figure 1. A.** Median upstream and downstream distance thresholds for the identification of DREs using standard proximity annotation or GREAT for the mouse (mm10) and human (hg19). **B.** Violin plot of upper distance limits for the identification of DREs using standard proximity annotation or GREAT. The percentage of genes with a distance threshold >100 kb is presented. **C.** Schematic overview of ICE-A, a tool for chromatin interaction-based annotation of genomic regions. **D.** Schematic representation of different DRE to target gene annotation methods, including standards proximity annotation, GREAT, and ICE-A interaction-based annotation. **E.** Percentage of candidate regions from Fulco *et al.* with H3K27ac, grouped based on if a significant effect on target gene expression was observed. **D-F.** Benchmarking of ICE-A using essential enhancers for cancer fitness from Chen et al. 2022. For interaction-based annotation approaches H3K4me3 PLAC-seq data (bin size 10kb) from HCT-116 and A549 cell lines are used, **D.** Schematic illustration of the different annotation methods including Proximity annotation, GREAT and ICE-A. In addition, ICE-A without proximity annotation is included. **E.** Fraction of essential enhancers in HCT-116 cell line identified with different annotation methods, presented as relative fraction compared to proximity annotation. **F.** Relative fraction of essential genes among the assigned target genes for essential enhancers.

S2

A

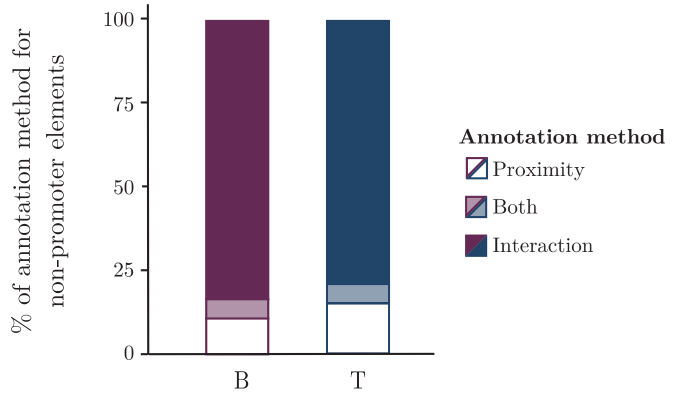


B

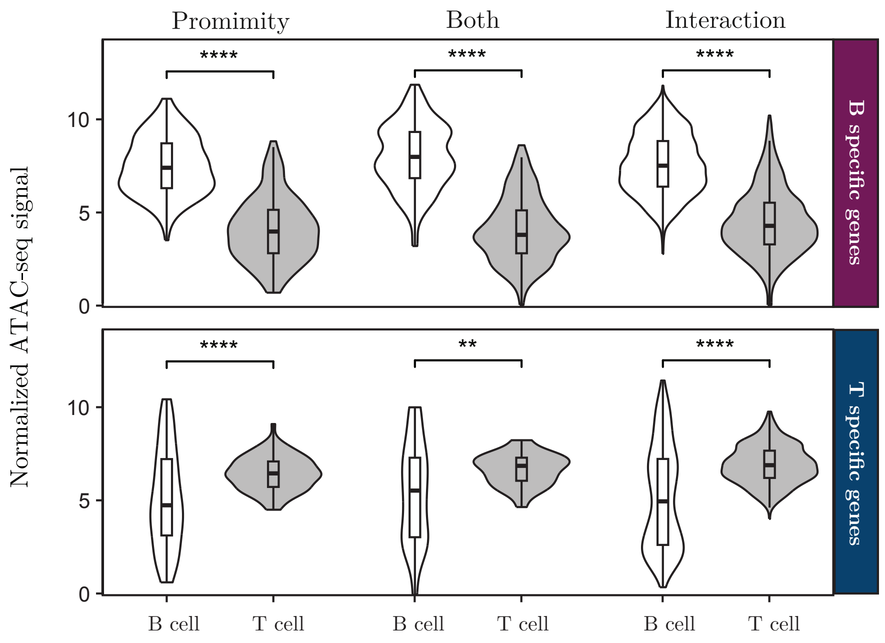


**Supplementary Figure 2. A.** Percentage of non-promoter elements associated to B- and T-cell specific genes identified using proximity or interaction-based annotation with ICE-A, related to Figure 2B. **B.** Chromatin accessibility of elements from A, in 230-238 B progenitor cell line and Scid.adh.2C2 T progenitor cell line. Statistical analyses are based on the Mann–Whitney *U*-test, **$p<0.01$, ****$p < 0.0001$.

S3

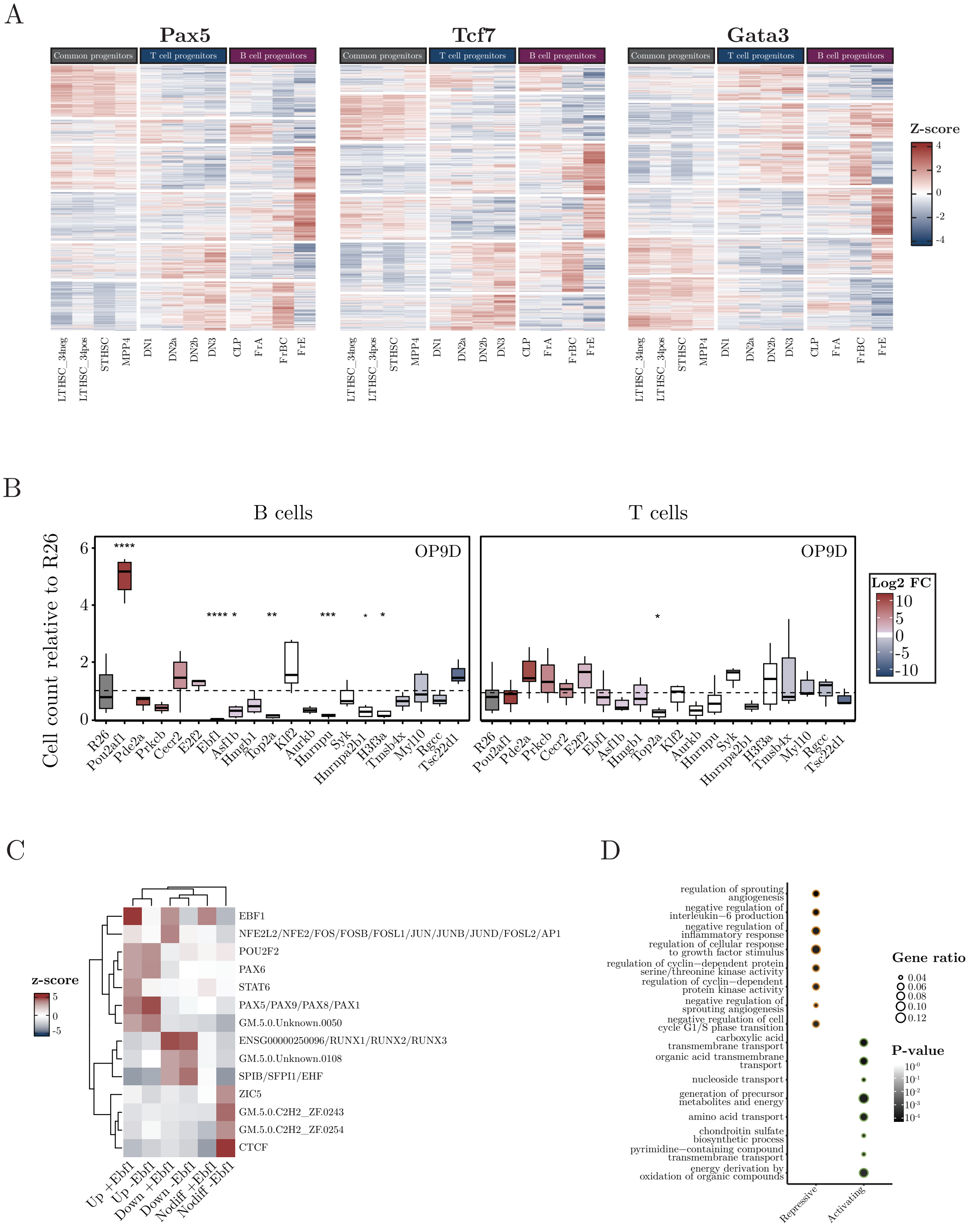
A
Pax5
Tcf7
Gata3
Common progenitors
T cell progenitors
B cell progenitors
Z-score
LTHSC_34neg
LTHSC_34pos
STHSC
MPP4
DN1
DN2a
DN2b
DN3
CLP
FrA
FrBC
FrE
B
B cells
T cells
OP9D
Cell count relative to R26
Log2 FC
R26
Pou2af1
Pde2a
Prkcb
Cecr2
E2f2
Ebf1
Asf1b
Hmgb1
Top2a
Klf2
Aurkb
Hnrnpu
Syk
Hnrnpa2b1
H3f3a
Tmsb4x
Myl10
Rgcc
Tsc22d1
C
z-score
EBF1
NFE2L2/NFE2/FOS/FOSB/FOSL1/JUN/JUNB/JUND/FOSL2/AP1
POU2F2
PAX6
STAT6
PAX5/PAX9/PAX8/PAX1
GM.5.0.Unknown.0050
ENSG00000250096/RUNX1/RUNX2/RUNX3
GM.5.0.Unknown.0108
SPIB/SFPI1/EHF
ZIC5
GM.5.0.C2H2_ZF.0243
GM.5.0.C2H2_ZF.0254
CTCF
Up +Ebf1
Up -Ebf1
Down +Ebf1
Down -Ebf1
Nodiff +Ebf1
Nodiff -Ebf1
D
regulation of sprouting angiogenesis
negative regulation of interleukin−6 production
negative regulation of inflammatory response
regulation of cellular response to growth factor stimulus
regulation of cyclin−dependent protein serine/threonine kinase activity
regulation of cyclin−dependent protein kinase activity
negative regulation of sprouting angiogenesis
negative regulation of cell cycle G1/S phase transition
carboxylic acid transmembrane transport
organic acid transmembrane transport
nucleoside transport
generation of precursor metabolites and energy
amino acid transport
chondroitin sulfate biosynthetic process
pyrimidine−containing compound transmembrane transport
energy derivation by oxidation of organic compounds
Repressive
Activating
Gene ratio
0.04
0.06
0.08
0.10
0.12
P-value

**Supplementary Figure 3. A.** Heatmap with the z-score of normalized gene expression in early lymphoid progenitors from the Immgen dataset (GSE100738), for genes bound by the lineage-specific TFs PAX5, TCF7 and GATA3. **B.** Diagram displaying the relative number of generated B- and T-cells compared to R26 control after 11 days of in vitro incubation of $KIT^{+}$ cells on OP9D stromal cells. The color represents log2FC *Wt* vs *Ebf1-/-* from RNA seq data. **C.** Comparative motif enrichment of a consensus set of open chromatin element from W*t* and *Ebf1*$^{-/-}$ ATAC-seq data, grouped by EBF1 occupancy and differential chromatin accessibility in W*t* vs *Ebf1*$^{-/-}$ (from Figure 3D). **D.** Gene ontology (GO) analysis of enriched biologic processes for genes with EBF1-addicted activation or repression. The top 50 genes based on the fold-changes in EBF1 degradation are included in the enrichment analysis.
